# Torsin ATPases are required to complete nuclear pore complex biogenesis in interphase

**DOI:** 10.1101/821835

**Authors:** Anthony J. Rampello, Ethan Laudermilch, Nidhi Vishnoi, Sarah M. Prohet, Lin Shao, Chenguang Zhao, C. Patrick Lusk, Christian Schlieker

## Abstract

Nuclear envelope herniations (blebs) containing FG-nucleoporins and ubiquitin are the phenotypic hallmark of Torsin ATPase manipulation. Both the dynamics of blebbing and the connection to nuclear pore biogenesis remain poorly understood. We employ a proteomics-based approach to identify MLF2 as a luminal component of the bleb. Using an MLF2-based live cell imaging platform, we demonstrate that NE blebbing occurs rapidly and synchronously immediately after nuclear envelope reformation during mitosis. Bleb formation is independent of ubiquitin conjugation within the bleb, but strictly dependent on POM121, a transmembrane nucleoporin essential for interphase nuclear pore biogenesis. Nup358, a late marker for interphase nuclear pore complex (NPC) biogenesis, is underrepresented relative to FG nucleoporins in nuclear envelopes of Torsin-deficient cells. The kinetics of bleb formation, its dependence on POM121, and a reduction of mature NPCs in Torsin deficient cells lead us to conclude that the hallmark phenotype of Torsin manipulation represents the accumulation of stalled NPC assembly intermediates.

## Introduction

Torsin ATPases (Torsins) are widely conserved proteins in metazoans and have essential, yet poorly understood roles. While Torsins are phylogenetically related to the well-characterized Clp/HSP100 proteins (Rose et al., 2015), they deviate from these ATPases in several fundamental aspects. Torsins are the sole members of the AAA+ ATPase superfamily to reside in both the lumen of the endoplasmic reticulum (ER) and the nuclear envelope (NE) (Laudermilch et al., 2016). Another unusual feature is that Torsins are inactive in isolation and require one of two membrane-spanning cofactors, LAP1 or LULL1, for ATPase activity (Zhao et al., 2013). This activation relies on a classical active site complementation mechanism, in which the luminal domain of LAP1 or LULL1 contribute an arginine finger that is notably absent from Torsin ATPases (Brown et al., 2014; Sosa et al., 2014). Apart from activating Torsins, these cofactors also modulate the oligomeric state of the Torsin assembly (Chase et al., 2017b). A steadily increasing number of mutations affecting this delicate assembly have been identified as causal factors in human pathologies. Some of these mutations destabilize essential intersubunit interactions at the Torsin-cofactor interface. Notably, this is the case for the highly debilitating movement disorder DYT1 dystonia (Brown et al., 2014; Demircioglu et al., 2016) where TorsinA was originally identified through a positional cloning approach (Ozelius et al., 1997). More recently, a LAP1 mutation was identified that severely limits the lifespan of affected individuals who suffer from diverse symptoms including dystonia and myopathy (Fichtman et al., 2019).

While the diverse set of Torsins exhibit tissue-specific expression (Jungwirth et al., 2010) and differential abilities to be stimulated by their distinctively localizing cofactors (Zhao et al., 2013), the shared hallmark phenotype that is observed upon their genetic manipulation from nematodes (VanGompel et al., 2015) to *Drosophila melanogaster* (Jokhi et al., 2013), mouse models (Goodchild et al., 2005; Liang et al., 2014; Tanabe et al., 2016), and tissue culture cells (Laudermilch et al., 2016; Naismith et al., 2004; Rose et al., 2014) is NE blebbing (Laudermilch and Schlieker, 2016). Major obstacles towards understanding Torsin function in this phenotypic context are the genetic redundancy between Torsin homologs in human tissue culture cells and mouse models (Kim et al., 2010; Laudermilch et al., 2016) and the essential nature of Torsins (Goodchild et al., 2005).

We previously presented a system that resolves both of these limitations by generating a quadruple Torsin deletion HeLa cell line (designated 4TorKO) in which all four Torsin genes (TorsinA, TorsinB, Torsin 2A and Torsin3A) have been deleted using CRISPR/Cas9 genome engineering. This 4TorKO cell line abundantly exhibits the hallmark cellular phenotype of NE blebbing in which the inner nuclear membrane (INM) bulges into the perinuclear space (PNS) to form an omega-shaped herniation. Ubiquitin (Ub) conjugates of the K48 linkage type are enriched in the lumen of the bleb in 4TorKO cells and in mouse models of Torsin dysfunction (Pappas et al., 2018). At the base of a bleb there is electron density with a uniform diameter and dimensions similar to the nuclear pore complex (NPC). This density can be decorated via immunogold labeling using Mab414 antibodies, which recognize several FG-rich NPC components termed FG nucleoporins (FG-Nups) (Laudermilch et al., 2016).

Whether a causal relationship exists between these NPC markers and bleb formation is largely unknown. However, the finding that nuclear transport is perturbed in *Caenorhabditis elegans* upon mutation of the TorsinA homolog OOC-5 (VanGompel et al., 2015) as well as the observation of altered *in situ* distribution of nuclear transport machinery in brain tissue of mouse models of dystonia (Pappas et al., 2018) further support a functional connection between Torsins and the NPC. Clearly, more insight into the molecular composition of these Nup-containing densities and their provenance is required to distinguish whether they are mature NPCs, products of stalled NPC biogenesis, or a result of NPC instability. One hurdle in testing kinetically resolved roles for Torsins in NPC biogenesis or homeostasis is the absence of bleb-specific live cell imaging markers. A better functional assignment for Torsins is additionally confounded by a lack of quantitative information about NPC number and assembly state in relation to bleb formation.

In this study, we quantify NPCs and observe a considerable reduction of mature NPCs with a concomitant increase of Nup-containing blebs in 4TorKO cells relative to wild type (WT) cells. These structures form in a strictly cell cycle-dependent fashion. We find that the protein Myeloid Leukemia Factor 2 (MLF2) is highly enriched in the lumen of newly forming blebs, allowing us to develop MLF2 derivatives as bleb-specific probes with broad utility for live cell imaging and functional characterization. Notably, bleb formation occurs rapidly and synchronously immediately following NE reformation after mitosis, a timing that is reminiscent of interphase NPC biogenesis. This dynamic buildup can be selectively perturbed by depletion of POM121, a Nup that is essential for interphase NPC biogenesis. These observations, as well as the diagnostic underrepresentation of the late NPC biogenesis marker Nup358 from Nup-containing blebs, establish a role for Torsins during interphase NPC biogenesis.

## Results

### Torsin-deficient cells exhibit reduced numbers of mature nuclear pores

To further explore a functional relationship between Torsin and NPCs, we first asked whether Torsin deletion leads to a reduction in the number of nuclear pores in asynchronously growing cells. To this end, we exploited our previously reported HeLa-based 4TorKO cell line and processed cells for standard transmission electron microscopy (EM) along with isogenic WT cells. As expected, WT cells featured an evenly spaced INM and ONM, with an average of 15.2 nuclear pores per 30 µM of NE (Fig. 1A, lower panel and Fig. 1B). As reported previously (Laudermilch et al., 2016), the blebbing phenotype was highly penetrant in 4TorKO cells (Fig. 1A, upper panel). The number of mature nuclear pores was significantly decreased relative to WT cells with an average of 11.2 nuclear pores per 30 µm of NE (Fig. 1B). This was accompanied by a corresponding increase of fuzzy electron density at the base of many blebs (Fig. 1B) that we have shown to represent FG-Nup containing assemblies (Laudermilch et al., 2016). Thus, the observed reduction of the number of nuclear pores with a concomitant increase of FG-Nup assemblies at the bases of blebs represents a highly robust phenotype.

**Figure 1:**
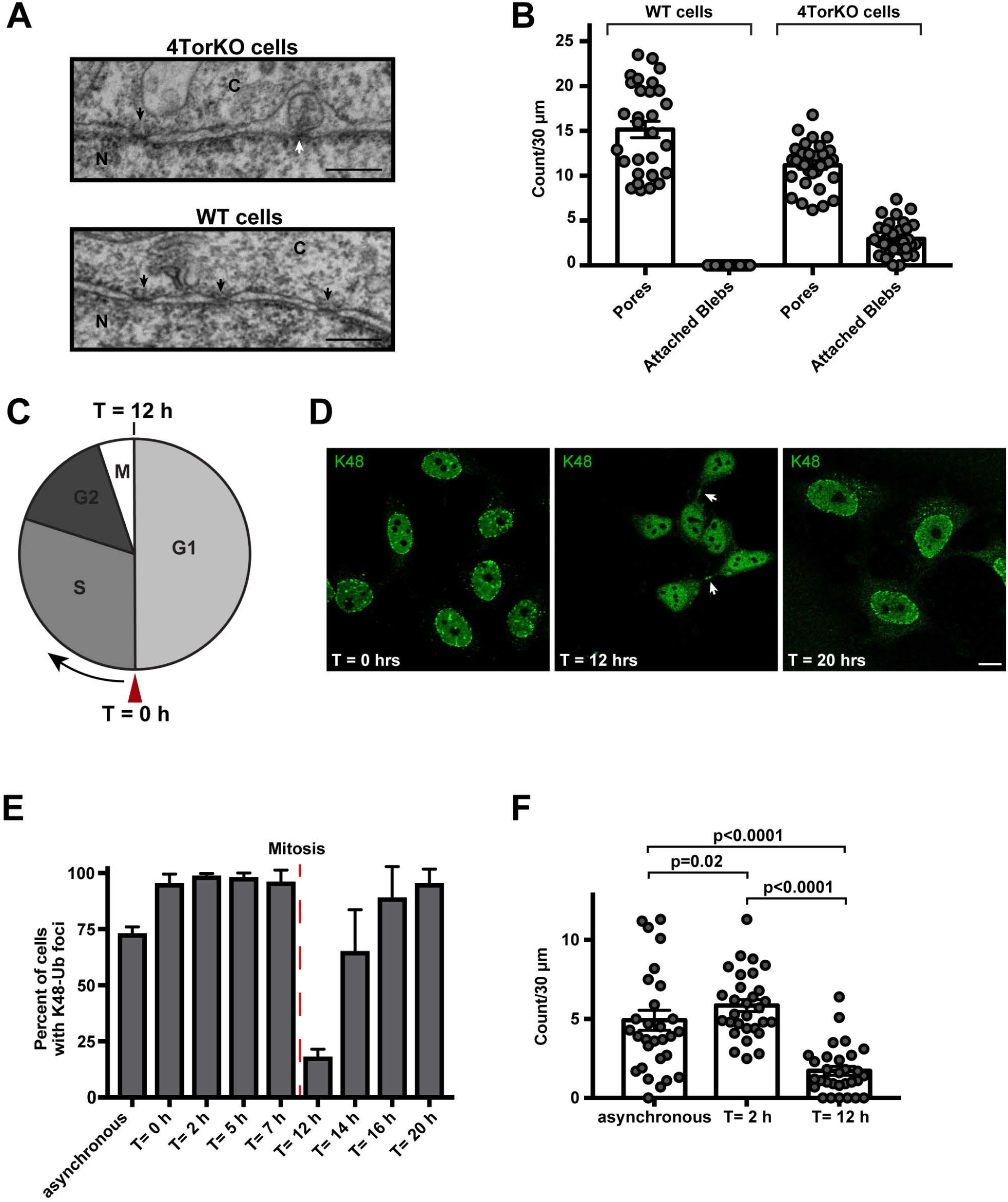
Nuclear envelope defects appear in a cell cycle specific manner in Torsin-deficient cells. (A) EM images of NE herniations in Torsin-deficient (4TorKO) HeLa cells. EM images of mature NPCs from wild type (WT) cells are shown for comparison. Black arrow: mature NPCs, white arrow: NE herniation, N: nucleus, C: cytoplasm. Scale bars are 250 nm. (B) Graph comparing the number of mature pores and NE herniations with visible neck regions found WT and 4TorKO cells. Quantifications are based on the number of mature pores or attached herniations observed per 30 µM of NE with each point on the graph representing an individual cross section. The average number of blebs per 30 µM of NE and the standard error of the mean are indicated by the bar graph and error bars, respectively. At least 30 cells were counted for each sample. (C) Diagram depicting the cell cycle stages in HeLa cells (∼20 h per complete cycle). In synchronization experiments, cells were arrested at the G1 to S phase transition (red arrowhead), which was designated as the 0-hour time point (T = 0 h). In D-F, the times given are the amount of time post-release from the double thymidine block. (D) Confocal images of three different time points from synchronized 4TorKO cells stained with a K48-Ub antibody. White arrow: midbodies. Representative scale bar is 10 µm. (E) Graph showing the percent of cells exhibiting K48-Ub foci at eight different time points from synchronized cells and an asynchronous control. Mitosis occurs between the 7- and 12-hour time points (dashed red line). Each time point shows the average of three independent experiments with at least 50 cells each. (F) Graph of the number of blebs observed per EM cross section in samples from synchronized cells at two time points or asynchronous cells. Each point on the graph represents the number of blebs per 30 µM of NE. The average number of blebs per 30 µM of NE and the standard error of the mean are indicated by the bar graph and error bars, respectively. At least 30 cells were counted for each sample.

### Nuclear envelope herniations form in interphase

A second important criterion supporting a connection to NPC assembly would be a possible cell cycle dependency of bleb formation. The latter assertion is based on prior work that cumulatively suggests that there are likely two biochemically (Doucet et al., 2010) and morphologically distinct mechanisms of NPC assembly, one occurring during post-mitotic NE reformation and the other during interphase (Otsuka and Ellenberg, 2018)(Fig. S1). In the latter case, there is an emerging consensus that NPC assembly begins from the inside of the nucleus on the INM (Doucet et al., 2010) likely through an inside-out evagination of the INM, which ultimately leads to fusion with the ONM (Otsuka et al., 2016). Interphase assembly might be under the control of cell cycle (Talamas and Hetzer, 2011) and other (McCloskey et al., 2018) regulators with a potential burst of this assembly mechanism in early G1 (Doucet et al., 2010; Otsuka et al., 2016; Weberruss and Antonin, 2016). If the Torsin knockout phenotype does indeed represent a stalling in NPC biogenesis, we would expect to observe the first signs of bleb formation early in G1. To investigate this possibility, we synchronized 4TorKO cells in early S-phase using the double thymidine block method (Bostock et al., 1971) and processed the cells for immunofluorescence at various times after release from the thymidine block (Fig. 1C, D). Anti-K48-Ub antibodies were used to score for bleb formation since K48-linked Ub is strongly enriched in the bleb lumen both in 4TorKO cells (Laudermilch et al., 2016) and in mouse models of DYT1 dystonia (Pappas et al., 2018). While the majority of asynchronously growing 4TorKO cells exhibited K48-Ub foci diagnostic of NE blebs (Fig. 1E), a striking cell cycle dependency was observed in synchronized cells. At the G1/S boundary (T = 0 h upon release of block), nearly all cells contain K48-Ub foci (Fig. 1D, E). Additionally, we observed essentially the same abundance for subsequent time points during S phase and early G2. Following mitosis, however, this number drops substantially (Fig. 1D, E). It should be noted that in our experience, the entire HeLa cell population is not sharply synchronized under conditions of the double thymidine synchronization. Some deviation from complete synchrony certainly exists (we estimate that about 85% of the population is well-synchronized). However, midbodies (that are also labeled with anti-K48-Ub, see middle panel in Fig. 1D) can be used as convenient diagnostic markers to assign cells to the time of late cytokinesis. Notably, K48-Ub NE foci are completely absent from the nascent NE of these dividing or recently divided cells. As cells proceed through G1, the number of K48-Ub foci steadily increases again and reaches a maximum at approximately 20 h post-release from the thymidine block (Fig. 1E), which is about one complete cell cycle. To directly confirm that the reduction of K48-Ub NE foci coincides with a loss of blebs, we also performed an analogous double thymidine block experiment and processed synchronized 4TorKO cells for EM at T = 2 h and T = 12 h post-release. These correspond to the time points of highest and lowest abundance of K48 foci, respectively. We observed that the number of blebs per cross section for each time point is in good agreement with the numbers derived from immunofluorescence (cf. Fig. 1E and F). We therefore conclude that the formation of blebs is a cell cycle-dependent process and that the majority of blebs are formed during G1.

### MLF2 is highly enriched in nuclear envelopes of 4TorKO cells

While the observed timing of bleb formation is consistent with a possible role for Torsins in NPC biogenesis (D’Angelo et al., 2006; Dultz and Ellenberg, 2010; Otsuka et al., 2016), a live cell imaging readout for bleb formation would enable higher temporal resolution. Moreover, live cell observations on a single cell level allow for a direct visualization of mitotic events and should resolve the issue of incomplete synchronization that limited the accuracy of our former measurements (cf. Fig. 1).

We therefore set out to identify a suitable marker that is specific to NE blebs. To this end, we utilized a comparative proteomics approach in which we compared the respective NE proteomes of 4TorKO verse WT cells. Briefly, we gently homogenized 4TorKO and WT cells and isolated NEs by a series of consecutive centrifugation steps (see Materials and Methods)(Fig. 2A). Since at least a subset of bleb luminal components appear to be conjugated to K48-Ub (Laudermilch et al., 2016), we solubilized NEs with mild detergent and immunoprecipitated the obtained extracts with K48-Ub-specific antibodies. Note that both the bleb structure and their reactivity to K48-Ub antibodies are preserved under these gentle NE isolation procedures (Fig 2A, inset) prior to addition of detergent. The resulting immunoprecipitates (IPs) were then resolved by SDS-PAGE and subsequently analyzed by mass spectrometry (MS).

**Figure 2:**
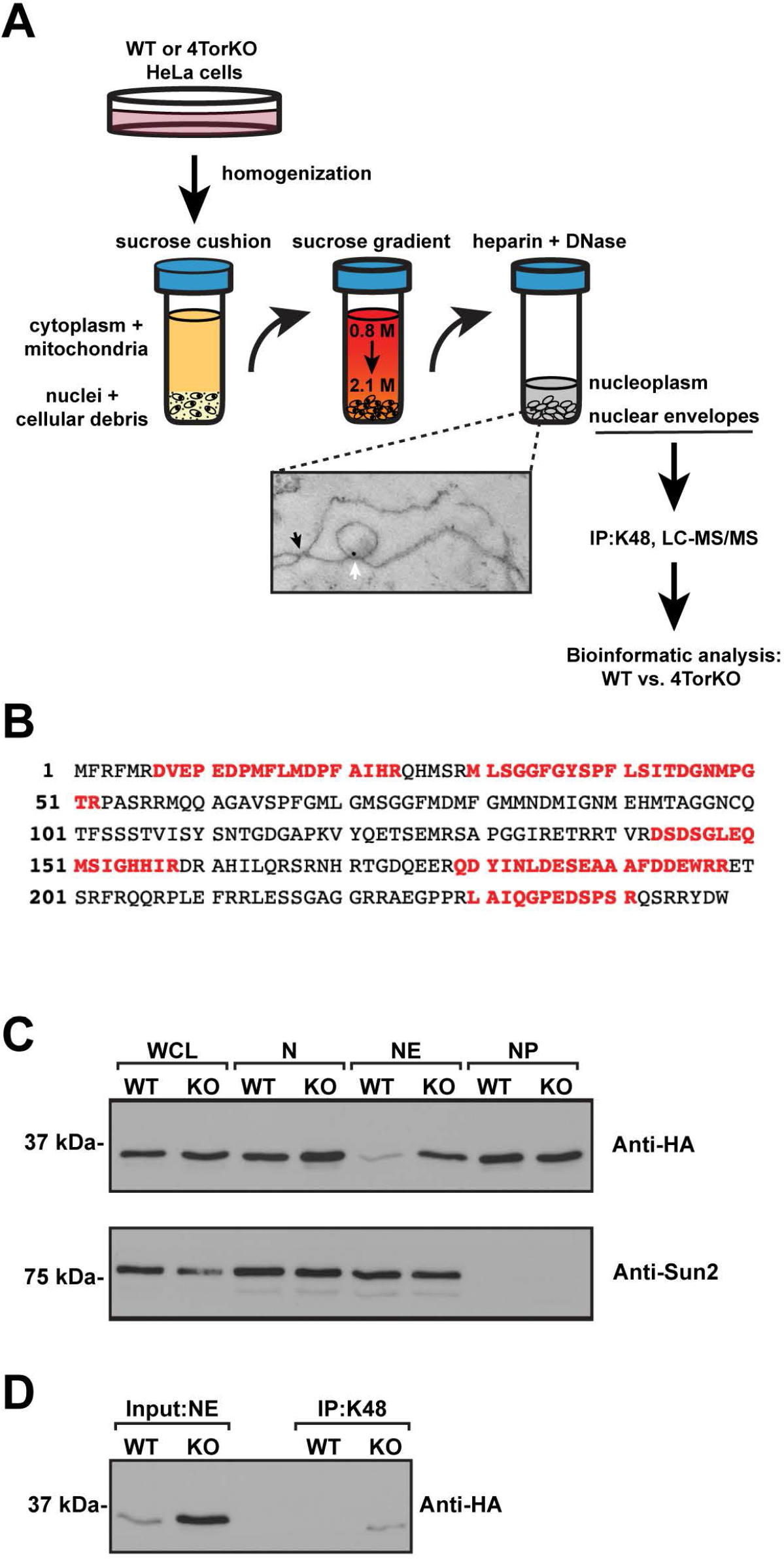
Identification of MLF2 as a molecular component of nuclear herniations in Torsin-deficient cells. (A) Overview of the subcellular fractionation and K48-Ub immunoprecipitation (IP) workflow. K48-Ub-linked candidate proteins were selected based on their fold-enrichment in 4TorKO cells over WT cells. Electron micrograph of NEs isolated from 4TorKO cells with immunogold labeling for K48-Ub is shown. Black arrow: NPC, white arrow: NE herniation containing K48-Ub. (B) Amino acid sequence of MLF2. A 36% sequence coverage was obtained by MS and the identified peptides are highlighted in red. (C) Western blot analysis of the subcellular fractions isolated from the workflow presented in (A) utilizing WT and 4TorKO cells stably expressing MLF2 with an endogenous C-terminal 3xHA tag. SUN2 was used as a NE marker. (D) Validation of the MLF2 association with K48-Ub chains in 4TorKO cells by co-IP. WT; wild type, KO; Torsin-deficient cells, WCL; whole cell lysate, N; nuclear fraction, NE; nuclear envelope fraction, and NP; Nucleoplasmic fraction.

One protein that stood out immediately was Myeloid Leukemia Factor 2 (MLF2) as it was identified with 36% sequence coverage in the 4TorKO sample compared to only 12% in the WT control sample (Table S1, Fig. 2B, identified peptides are highlighted in red). To confirm this enrichment, we employed CRISPR/Cas9 genome engineering and installed a C-terminal tandem HA tag on MLF2 at the endogenous locus in both WT and 4TorKO genetic backgrounds. Clonal cell lines were isolated from WT and 4TorKO backgrounds with equivalent MLF2-HA expression levels (cf. whole cell lysates, WCL, Fig. 2C). Using these cell lines, we conducted subcellular fractionations to analyze the relative amount of MLF2 in nuclear, NE, and nucleoplasmic fractions. After fractionation, the corresponding samples were solubilized in SDS and subjected to SDS-PAGE and immunoblotting. While MLF2-HA levels were approximately the same in whole cell lysates, we observed slightly more MLF2 in the nuclear fraction of 4TorKO cells (Fig. 2C). A further deconvolution of nuclei into NEs and nucleoplasm revealed a major enrichment of MLF2 in the NEs of 4TorKO cells relative to WT cells, while the levels were comparable in the nucleoplasmic fraction (Fig. 2C). The INM protein Sun2 was additionally monitored via immunoblotting to confirm successful fractionation (Tsai et al., 2019). Finally, we subjected detergent extracts of both NE fractions to immunoprecipitation with anti-K48-Ub antibodies followed by SDS-PAGE and immunoblotting. In line with our original MS-based experiment, we detected MLF2 in the IPs from 4TorKO NEs but not from WT NEs (Fig. 2D). These results led us to explore the potential use of MLF2 as a live cell imaging tool to investigate the potential role of Torsin ATPases in NPC biogenesis.

### Establishing MLF2 as a bleb-specific marker

To begin, we engineered an MLF2-GFP fusion protein, and asked whether MLF2 localizes to blebs. 4TorKO cells were transfected with MLF2-GFP and we then monitored the localization of the fusion protein relative to blebs (direct fluorescence versus anti-K48-Ub signal) by immunofluorescence. We observed a punctate pattern for MLF2-GFP at the nuclear periphery combined with a diffuse nucleoplasmic staining (Fig. 3A). These observations are in excellent agreement with our biochemical fractionation (Fig. 2C). More importantly, we observed a correlation in 4TorKO cells displaying both K48-Ub and MLF2 foci (Fig. S2A-C), and a considerable, though not complete, degree of colocalization between K48-Ub staining and MLF2-GFP, which is also apparent in a line scan analysis (Fig. S3A).

**Figure 3:**
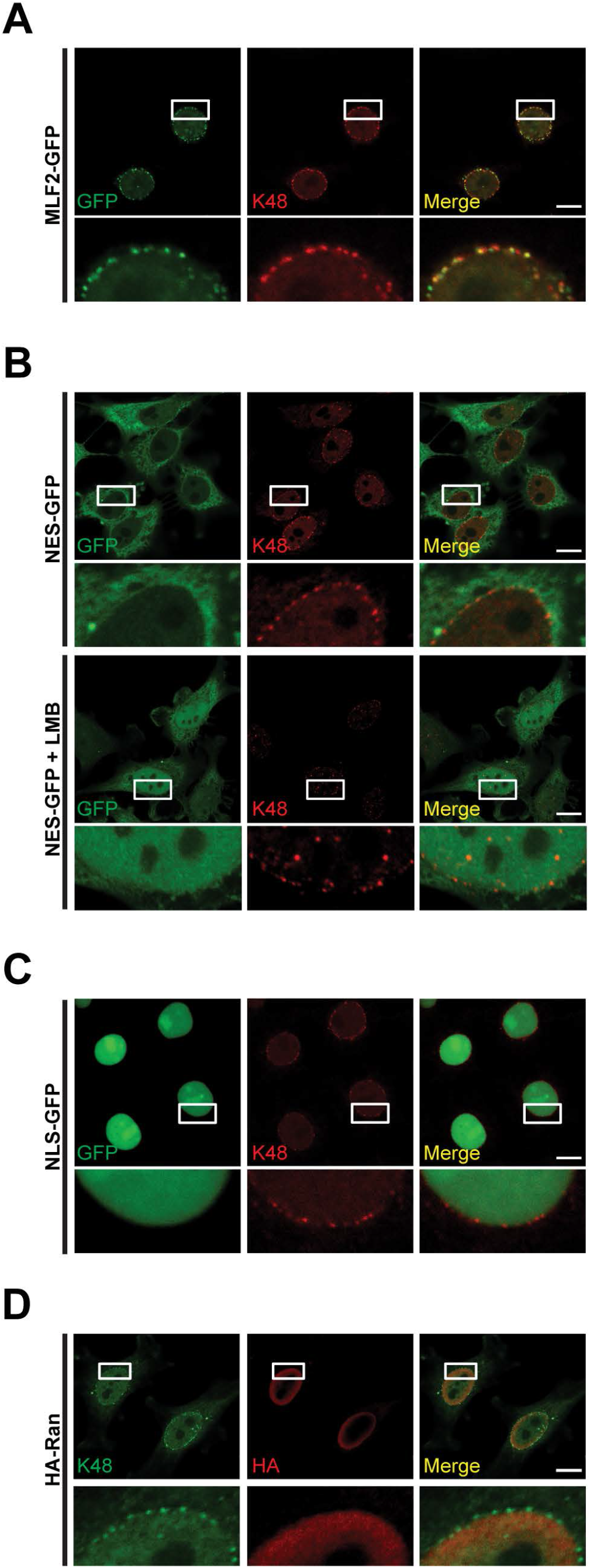
Nuclear herniations exhibit specificity in their molecular inventory. (A-C) Confocal microscopy of 4TorKO cells transfected with MLF2-GFP (A), NES-GFP (B), or NLS-GFP (C) and counterstained with anti-K48-Ub (red) to label NE herniations. (B) Nucleo-cytoplasmic transport competency was assessed through the treatment of NES-GFP transfected cells with 10 ng mL^-1^ Leptomycin B (LMB), a CRM1-dependant nuclear export inhibitor, for 4 hours prior to fixation. (D) Confocal images of Torsin-deficient cells transfected with HA-Ran (red) and counterstained with anti-K48-Ub (green). Representative scale bars are 10 µm.

The question arises whether the observed MLF2-GFP localization is indeed a specific indicator of NE bleb formation or if the perinuclear foci formation merely results from perturbed nuclear export. We therefore asked if GFP variants with a nuclear export signal (NES-GFP) or nuclear import signal (NLS-GFP) give rise to a similar or distinct localization pattern. Both NES-GFP and NLS-GFP variants showed the expected extranuclear and nucleoplasmic localization, respectively (Fig. 3B, upper panel, and Fig. 3C). To validate the functionality of NES-GFP and explore the nucleo-cytoplasmic transport competency of 4TorKO cells, we treated cells with Leptomycin B (LMB), a CRM1-dependant nuclear export inhibitor (Kudo et al., 1998). As expected, we observed a strong nucleoplasmic GFP signal upon inhibition of nucleo-cytoplasmic trafficking (Fig. 3B, lower panel), excluding the formal possibility that NES-GFP never enters the nucleus. Additionally, we did not observe an enrichment of HA-tagged Ran (Fig. 3D), a major player in nuclear transport (Adam et al., 1992; Lui and Huang, 2009). Thus, MLF2 is a highly specific marker for NE aberrations in 4TorKO cells.

We next asked whether MLF2-GFP distinctively localizes to NE blebs, a question that is best addressed via EM, which also provides information about membrane topology. We processed MLF2-GFP-expressing 4TorKO cells for EM via high pressure freezing and subjected sections to immunogold labeling with anti-GFP antibodies. 4TorKO cells displayed the typical accumulation of NE blebs (Laudermilch et al., 2016). We observed a striking enrichment of immunogold labeling of MLF2-GFP in these blebs and can clearly assign this accumulation to the bleb lumen that is enclosed by the INM (Fig. 4A). At a low frequency, we additionally observe cases in which gold particles concentrate in direct juxtaposition of a deformed INM (Fig. 4B, C). These could represent early bleb intermediates in which evaginations of the INM begin to form. The early addition of MLF2 to bleb intermediates would imply that MLF2 might be added before K48-Ub conjugation occurs. Supporting this idea, we observe more 4TorKO cells with MLF2 foci earlier than K48-Ub in G1 phase (Fig. 4D). We therefore conclude that MLF2 has the potential to be an effective tool in elucidating the dynamics of bleb formation.

**Figure 4:**
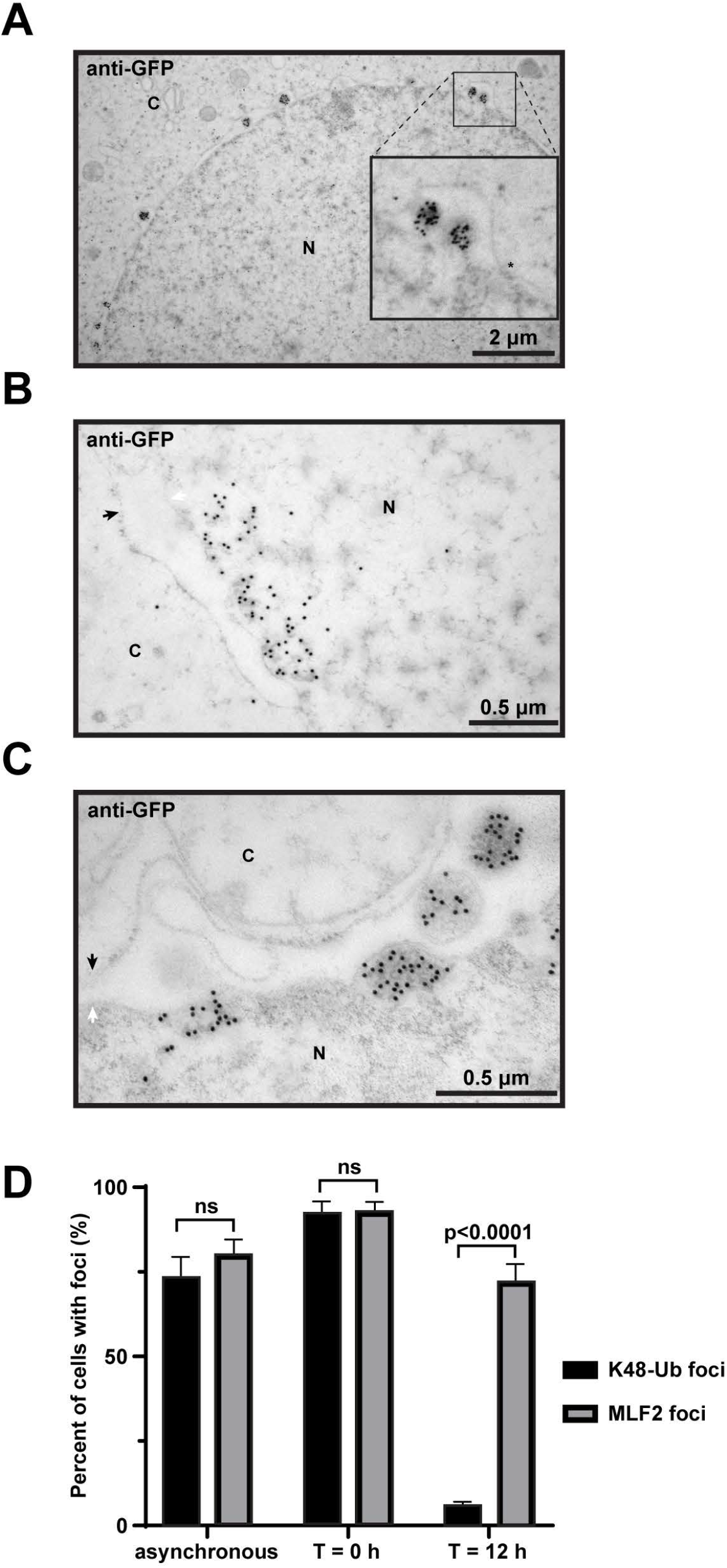
MLF2 is enriched in nuclear herniations. (A) Torsin-deficient cells were transfected with MLF2-GFP and analyzed by immunoelectron microscopy to visualize the subcellular localization of the GFP fusion proteins. Black arrow: outer nuclear membrane, white arrow: inner nuclear membrane, N: nucleus, C: cytoplasm, asterisk: NPC. (B and C) EM images showing MLF2-GFP enrichment at sites of increased membrane curvature on the inner nuclear membrane. (D) Percentage of cells showing MLF2-HA foci and K48-Ub foci from an asynchronous population, cells at the G1 to S phase transition (T = 0 h), and cells emerging from mitosis in early G1 (T = 12 h). Error bars indicate ± SD.

### Live cell imaging with MLF2-GFP reveals rapid and synchronous formation of nuclear envelope blebs

Resolving the dynamics of bleb formation relative to NE reformation during mitosis requires a non-invasive, robust live cell imaging platform. To this end, we generated a 4TorKO cell line stably expressing MLF2-GFP and mScarlet-Sec61β through retroviral transduction. To mitigate the potential occurrence of any artificial morphological effects on cells resulting from the constitutive activation of either gene, we employed a doxycycline (Dox)-inducible promoter system to control the expression levels of both genes. Furthermore, we utilized lattice light sheet microscopy (LLSM) as it provides rapid three-dimensional image acquisition with reduced photobleaching thus allowing the acquisition of longer time series data. Since we crudely assigned bleb formation to the early G1 phase (cf. Fig. 1E), we identified prometaphase/metaphase mitotic cells based on their round appearance and visible metaphase plates (Fig. 5A) and followed MLF2-GFP and mScarlet-Sec61β into early G1 (Supplemental Video 1). For standardization, the onset of anaphase was arbitrarily defined as T = 0 s. In agreement with previous observations, NE reformation after open mitosis occurred between 400-500 s after anaphase onset (Fig. 5A) (Dechat et al., 2004). At about 700 s, the first small MLF2-GFP foci appear in the nuclear periphery, while larger ones are observed at later times (Fig. 5A). Additionally, a small number of foci appear to form at some distance to the nuclear periphery in the nucleoplasm (Fig. 5A and Supplemental Video 1). Since we observed this trend repeatedly, we scrutinized this process further in an independent experiment to obtain a deconvoluted image series. We observed that the vast majority of these seemingly nucleoplasmic “outliers” are in fact closely associated with evaginations or wrinkles of the NE as judged by their colocalization with mScarlet-Sec61β (Fig. 5B, Supplemental Video 2).

**Figure 5:**
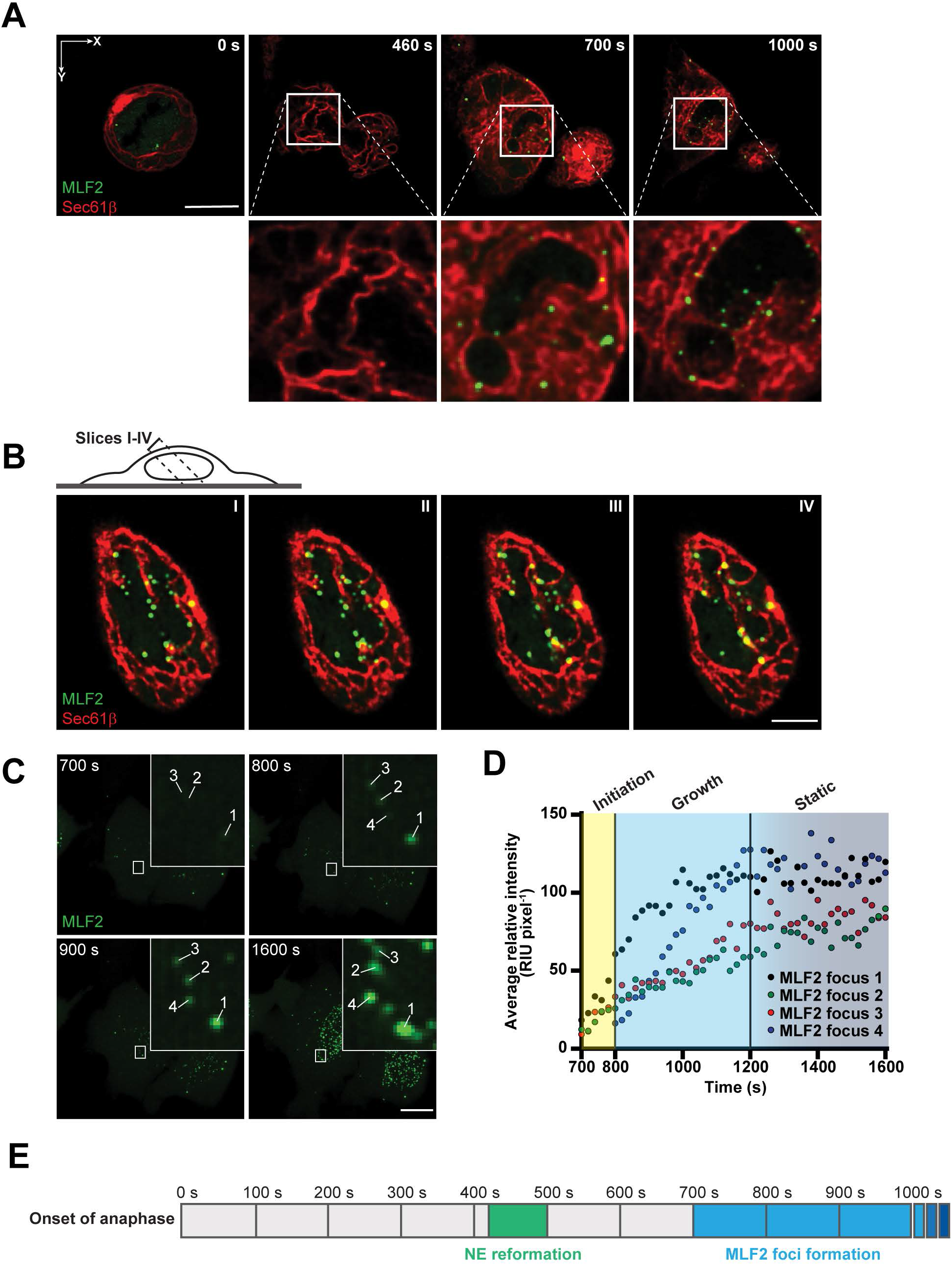
Utilizing MLF2 as a live cell imaging marker to visualize the cell cycle-dependent dynamics of NE blebbing. (A) Orthogonal sections of 4TorKO cells expressing mScarlet-Sec61β and MLF2-GFP under a Dox-inducible promoter. Sections are from four time points in a 3D data set from a 1600 s time series. Time 0 s was defined by the onset of anaphase and time 460 s shows the completion of NE reformation. NE herniations arise during a narrow window of early G1 phase immediately following cytokinesis (∼700 s) and persist through interphase. Data correspond to maximum intensity projections of 10 slices from a 3D image stack (z-axis) shown in Supplemental Video 1. (B) Sequential orthogonal sections highlighting the presence of MLF2-GFP foci on branched membrane networks connected to the NE. (C) Representative montage of the fluorescence intensity growth for four MLF2-GFP foci from the left daughter cell shown in panel A and Supplemental Video 1. Fluorescence intensities were measured from the initial appearance of foci to the end of image acquisition (700-1600 s). (D) Graphical representation depicting the increase in fluorescence intensity of MLF2 foci from (C). Times 700-800 s is designated as “Initiation phase”, times 801-1200 s is designated as “Growth phase”, and times 1201-1600 s is designated as “Static phase.” (F) Timeline displaying NE reformation (∼ 400-500 s), and the onset of herniation formation and growth (700+ s), which is represented by a blue gradient. The ranges of NE reformation and MLF2 foci formation were determined by observing five different cells (n = 5). Representative scale bars are 10 µm.

Since LLSM is superior to conventional fluorescence microscopy in terms of photobleaching, we were able to closely resolve the growth of individual foci over time. Focusing on the formation and maturation of a subset of MLF2-GFP foci in an individual daughter cell, we observed an initial steep growth phase that reached a maximum fluorescence intensity around 1200 s after anaphase onset (Fig. 5C-E). After this rapid growth phase, the foci appear to be static (Fig. 5D). Thus, the formation of the blebs is far more rapid than previously inferred from utilizing K48-Ub as a readout in fixed cells (cf. Fig. 1E). Another unexpected observation is the synchrony with which the bleb formation occurs (Fig. 5A, D, E, and Supplemental Video 1). Based on our time-resolved recordings, we estimate that the vast majority of initiation events can be narrowed down to a ∼100 s time window starting at ∼700 s after anaphase onset, which is in agreement with the appearance of NPC intermediates and subsequent formation of nascent NPCs from previous reports utilizing diverse microscopic methodologies (D’Angelo et al., 2006; Otsuka et al., 2016; Otsuka et al., 2018a).

### Ubiquitin conjugation is dispensable for bleb formation

Ubiquitin was the first-characterized marker to label NE blebs in a TorsinA-deficient mouse model (Liang et al., 2014). However, it has been unclear if a functional relationship between ubiquitylation and bleb formation exists. To test for a possible requirement, we asked whether we could engineer MLF2 to recruit Ub-modifying enzymatic activities to the bleb lumen. To this end, we engineered a construct consisting of a N-terminal MLF2 moiety fused to a deubiquitinating enzyme (DUB) domain derived from M48, the largest tegument protein of murine cytomegalovirus (Schlieker et al., 2005), followed by a C-terminal FLAG tag to create MLF2-M48^WT^ (Fig. 6A, B). This DUB domain potently deconjugates K48-linked Ub chains (Schlieker et al., 2007), the linkage type that is present in blebs (Laudermilch et al., 2016; Pappas et al., 2018) (Fig. 1D). As a control, we engineered a catalytically inactive variant in which the active site cysteine is mutated to an alanine, MLF2-M48^C23A^.

**Figure 6:**
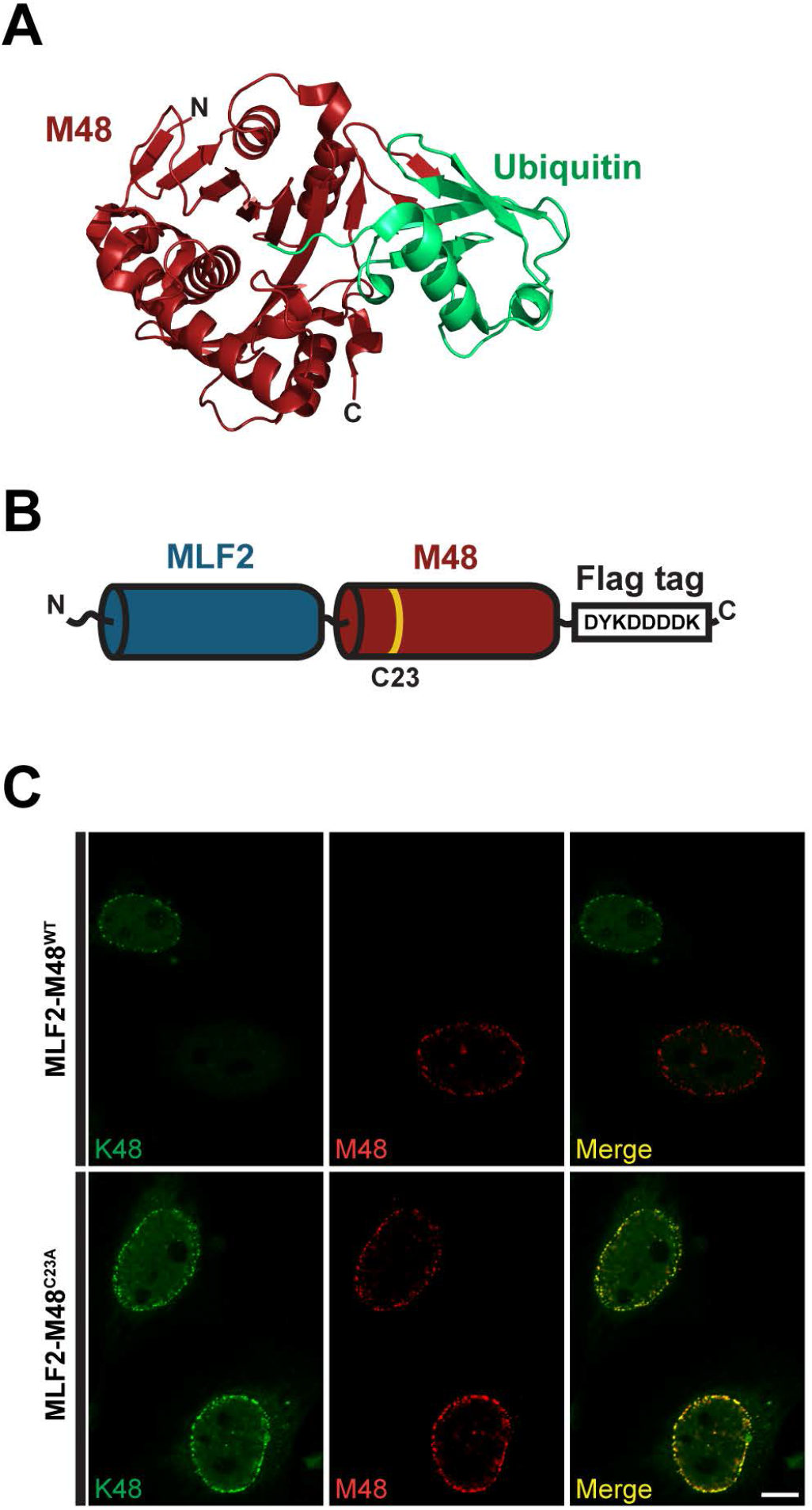
The formation of blebs in 4TOrKO cells is independent of K48-linked ubiquitin and precedes the formation of K48-Ub foci. (A) Structure of murine cytomegalovirus tegument protein M48 (red) bound to Ub (green) (PDB ID: 2J7Q). (B) Schematic representation of MLF2 with M48 fused to its C terminus (MLF2-M48^WT^). A C-terminal Flag tag is presenting the fusion construct for immunofluorescent analysis. The location of the catalytic cysteine residue is highlighted in yellow. (C) Representative confocal images of 4TorKO cells expressing MLF2-M48^WT^ or a catalytically inactive mutant MLF2-M48^DUB^-C23A. Cells were co-stained with anti-K48-Ub and anti-Flag antibodies. Representative scale bar is 10 µm.

4TorKO cells were transfected with either variant and processed for immunofluorescence with anti-FLAG and anti-K48-Ub antibodies 24 h post transfection. With a transfection efficiency of about 50%, non-transfected cells serve as a convenient control. While non-transfected cells display the expected K48-Ub foci phenotype, K48-Ub foci are virtually absent from cells expressing MLF2-M48^WT^ (Fig. 6C). However, MLF2-M48^WT^ is still found in perinuclear foci in a manner identical to canonical MLF2 staining (cf. Fig 6C and Fig. 3A or Fig. S2A), indicating that blebs form independently of K48-Ub enrichment. In the case of catalytically inactive MLF2-M48^C23A^, the signals of K48-Ub and MLF2-M48^C23A^ show the expected degree of colocalization, indicating that it is indeed the DUB activity that is responsible for the lack of K48-Ub signal in MLF2-M48^WT^-transfected cells.

These data argue against a critical role for K48-Ub conjugation in bleb formation, while establishing MLF2 as a useful tool to recruit specific enzymatic activities to NE blebs.

### Diagnostic absence of late NPC assembly markers relative to FG nucleoporins

Nup358 is a cytosolic-facing Nup that is recruited to a nascent NPC after the assembly of the bulk of the FG-Nups and the fusion of the INM and ONM (Otsuka et al., 2016). Therefore, as we previously proposed, the absence of Nup358 from FG-Nup containing blebs may provide a useful tool to assess whether these blebs are formed at sites of stalled NPC biogenesis (Chase et al., 2017a). We therefore imaged WT and 4TorKO cells via three-dimensional structured illumination microscopy (3D-SIM) and compared the localization of Nup358 and other FG-Nups using anti-Nup358 antibodies and the pan anti-FXFG antibody, Mab414, respectively. While we recognize that Mab414 is capable, in principal, of labeling Nup358 (Wu et al., 1995), it was the only antibody tested that provided the necessary specificity and signal-to-noise ratio to confidently assign NPCs using SIM. Moreover, it is established that Mab414 favors labeling Nup62 (Davis and Blobel, 1986) because it has more FXFG repeats and is found at higher copy numbers in the NPC (when fully formed) compared to Nup358 (Ori et al., 2013). Thus, the contribution of any Mab414-specific Nup358 labeling would likely be negligible.

Consistent with the idea that we can detect fully formed NPCs by SIM, we observe a near-complete colocalization of the Mab414 and Nup358 signals in WT cells in focal planes that illuminate the nuclear surface (Fig. 7A). Moreover, in mid-planes where NPCs are viewed by cross section, it is apparent that the Nup358 signal is spatially separated from the Mab414 signal, with the latter being more proximal and Nup358 being more distal relative to the nuclear interior. This is in agreement with our current understanding of NPC structure (Lin and Hoelz, 2019; Rout et al., 2000; Schwartz, 2016; von Appen et al., 2015). In this view, it is also clear that the Mab414 does not detectably label Nup358, confirming our prior assumption with respect to the specificity of Mab414. In contrast, in 4TorKO cells we observed irregularly shaped focal areas with a diameter of up to 5 µm in which we saw robust staining with Mab414 at a density that is comparable to WT cells, but with a notable absence of the anti-Nup358 label (Fig. 7A). This suggests that these areas may represent the accumulation of stalled intermediates during NPC assembly.

**Figure 7:**
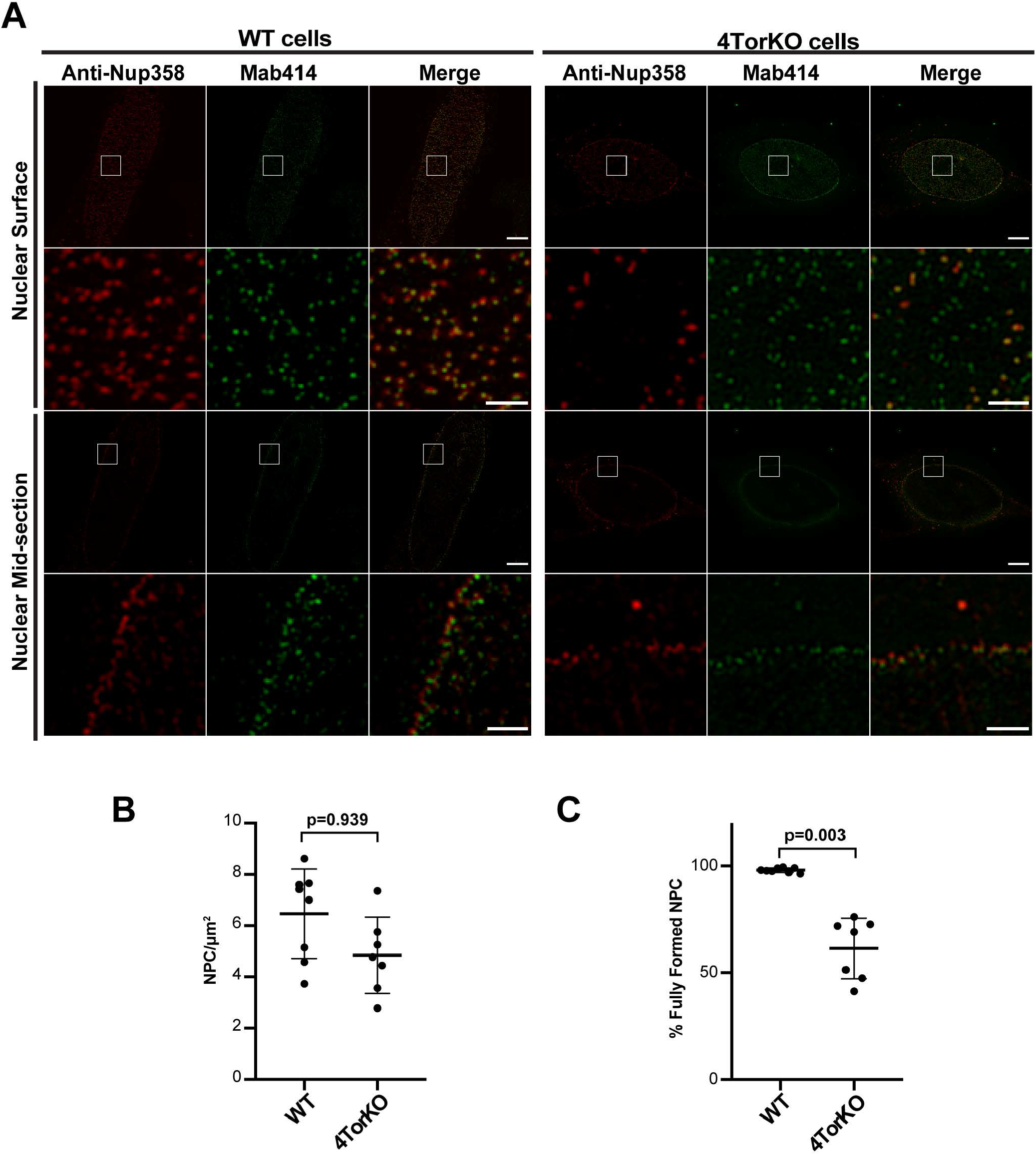
Torsin-deficient cells display defective assembly of NPCs. (A) 3D-SIM micrographs of WT and 4TorKO HeLa cells showing NPCs staining with anti-Nup358 (red) and Mab414 (green) antibodies from approximate nuclear surface and center. Scale bars are 5 µm (inset, 1 µm). (B) Plot showing the quantification of NPC density defined by dividing the number of Mab414 foci by the nuclear surface area from seven total cells (n = 7 from three independent experiments). (C) Plot showing the percentage of fully formed NPCs defined by the number of overlapping anti-Nup358 and Mab414 foci on the nuclear surface from seven cells each (n = 7 from three independent experiments). Error bars indicate ± SD.

Finally, we quantified the total number of Mab414-foci based on seven nuclei each (from three independent experiments) from WT and 4TorKO cells, which we interpret as the sum of mature and immature NPCs. We observe a modest, albeit insignificant, reduction in the density of Mab414 foci in 4TorKO cells suggesting that there is not a major reduction in NPCs or NPC biogenesis sites in the absence of Torsins (Fig. 7B). However, when we compared the fraction of colocalizing Nup358 and Mab414 foci as a measure for mature NPCs with the number of Mab414 sites arbitrarily expressed as 100%, we observed a ∼40% reduction in mature NPCs in 4TorKO cells relative to WT cells (Fig. 7C). This result is in good agreement with the observed reduction of NPCs and the concomitant increase of bleb-localized, FG Nup-containing densities in 4TorKO cells in electron micrographs (cf. Fig 7C and Fig. 1B).

In conclusion, the observed underrepresentation of Nup358 from sites containing FG Nups is consistent with the interpretation that a large proportion of FG Nup-containing Nup assemblies are devoid of cytoplasmic fibrils likely because NPC biogenesis is stalled at a step prior to INM/ONM fusion.

### POM121 is essential for bleb formation

Having shown that NE blebs in 4 TorKO cells feature NPC-like structures at their bases, a key question that remains is whether a causal relationship of NPC components for NE blebbing exists. An essential requirement of an NPC component for bleb biogenesis would lend significant credence to the idea that NE blebs represent “frozen intermediates” during NPC formation (Chase et al., 2017a; Laudermilch and Schlieker, 2016). Previous work established that the mitotic insertion of NPCs during open mitosis requires ELYS, while insertion of NPCs after reformation of the NE is independent of ELYS but highly sensitive to the depletion of the transmembrane Nup POM121 (Doucet et al., 2010; Franz et al., 2007). Having shown that NE blebs form *after* NE reformation (Fig. 5A), and given that these are topologically identical and morphologically similar to NPC biogenesis intermediates (Laudermilch et al., 2016; Otsuka et al., 2016), we reasoned that this selective dependency could be exploited by directly testing whether ELYS and POM121 are implicated in bleb formation.

As a first step, we depleted either ELYS or POM121 in 4TorKO cells via siRNA-mediated silencing and scored cells for any effects on K48-Ub foci formation. While both siRNA treatments potently reduced the RNA levels of their targets, only POM121 led to a stark reduction of K48-Ub focus formation, while silencing of ELYS had no significant effect (Fig. 8A-C). Since K48-Ub foci form relatively late during bleb formation (cf. Fig. 4D), we additionally monitored bleb formation via EM to directly visualize membrane deformation during bleb biogenesis under knockdown conditions (Fig. 8D and E). These results mirrored our observations using K48-Ub as readout. Bleb formation was essentially unperturbed in ELYS-silenced 4TorKO cells, whereas depletion of POM121 resulted in a stark, statistically significant decrease in the number of blebs per NE length (Fig. 8D). Together, these data establish an epistatic relationship between Torsins and the NPC component POM121. Considering that our knockdown approach did not completely eliminate POM121 on the transcript level (Fig. 8B) but nevertheless leads to a material reduction in bleb formation (Fig. 8D, E), it seems reasonable to deduce that POM121 is strictly required for bleb formation.

**Figure 8:**
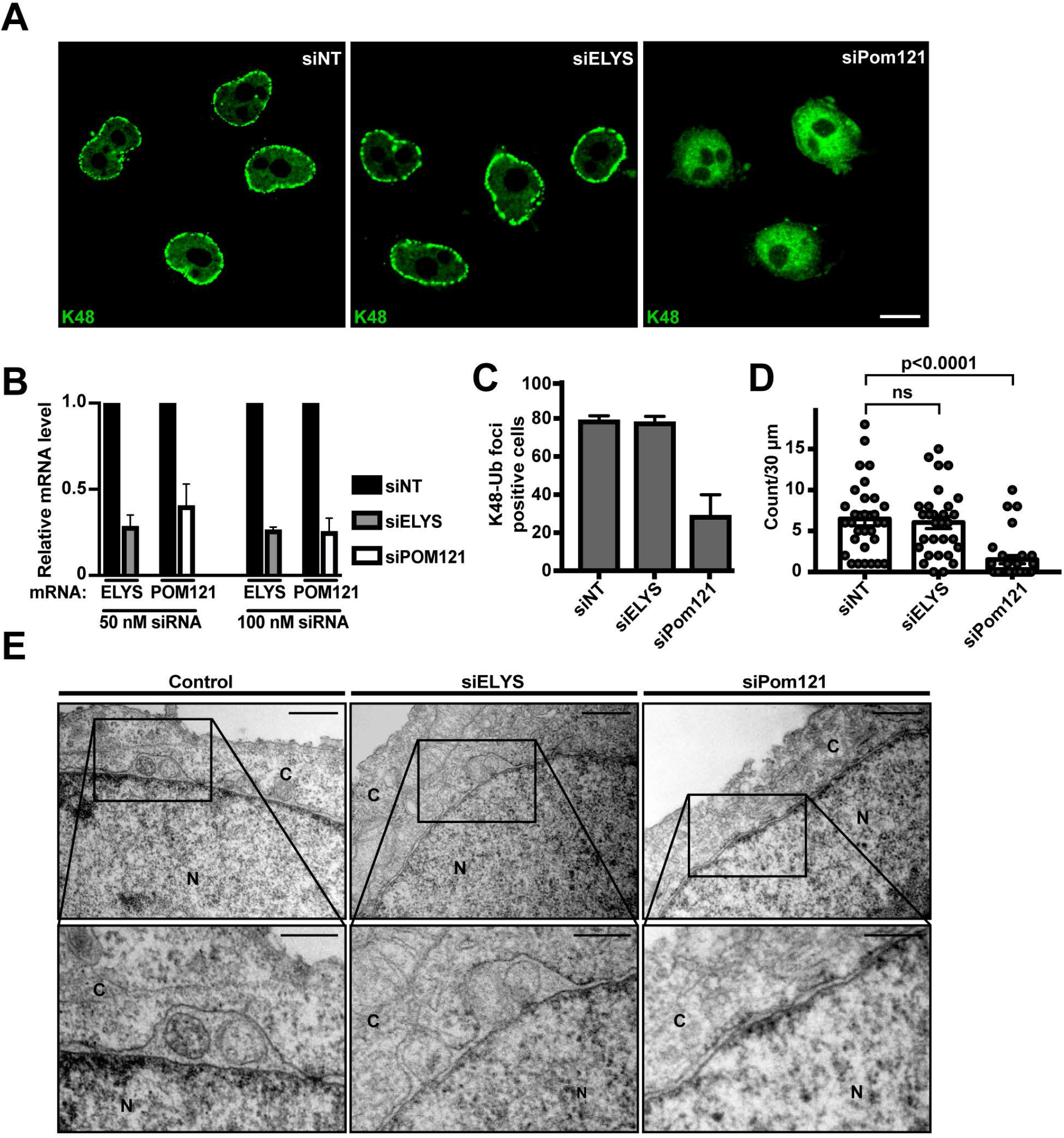
POM121 depletion relieves nuclear envelope abnormalities. (A) Confocal images of cells treated with 100 nM non-targeting (NT) siRNA, ELYS siRNA or POM121 siRNA. Cells were stained with an anti-K48-Ub antibody. Representative scale bar is 10 µm. (B) The relative amount of ELYS and POM121 mRNA transcripts following 50 nM or 100 nM siRNA treatments were analyzed by RT-qPCR with the indicated primers. Relative mRNA levels were normalized to the siNT control. The values are means of three independent replicates (n = 3) and error bars indicate ± SD. (C) Graph of the percent of cells with K48-Ub foci after treatment with 100 nM of the indicated siRNAs. Each result is the average of three independent experiments with at least 75 cells each. (D) Graph of the number of blebs observed per EM cross section upon treatment with 100 nM of the indicated siRNA. Each point shows the number of blebs per 30 µM of NE in an individual cross section. The average blebs per 30 µM of NE is shown by the bar graph and the standard error of the mean is shown in the error bars. At least 30 cells from each sample were counted. (E) EM images of 4TorKO control cells or cells treated with 100 nM siRNA targeting ELYS or Pom121. The bottom panel is an enlarged view of the boxed region in the top panel. N: nucleus, C: cytoplasm.

## Discussion

Nuclear envelope blebbing has been observed in developmentally regulated processes or upon genetic perturbation of Nups in numerous model organisms (Thaller and Lusk, 2018). Genetic ablation or mutation of specific Nups fall into the latter category, with *NUP116* deletions being examples of NE blebs with morphological similarities relative to the ones seen in 4TorKO cells (Onischenko et al., 2017; Wente and Blobel, 1993). The identification of a subset of FG-Nups at the electron-dense base of NE blebs in Torsin-deficient cells, as well as the finding that the diameter of this density is similar to mature NPCs (Laudermilch et al., 2016), previously suggested that Torsins could be implicated in NPC biogenesis (Chase et al., 2017a; Laudermilch and Schlieker, 2016). The recent discovery an inside-out evagination in the context of interphase NPC biogenesis (Otsuka et al., 2016) additionally revealed a phenomenon similar to the effects seen upon Torsin manipulation (Fig. 1A) (Chase et al., 2017a; Otsuka et al., 2018b; Weberruss and Antonin, 2016). Thus, several similarities exist between the two phenomena that relate Torsins to NPC biogenesis.

In this study, we asked whether a causal relationship can be established between Torsins and NPC biogenesis. We observed that the number of mature NPCs is strongly reduced in 4TorKO cells, with 23% of NPC-like structures being located at the base of NE blebs (Fig. 1A, B). Using Ub as a marker for blebs in the context of fixed cells, the timing of bleb formation falls mostly within the early G1 phase of the cell cycle (Fig. 1C-F), a window of when a burst of interphase NPC biogenesis has been observed (Dultz and Ellenberg, 2010). The observed penetrance of this Torsin knockout phenotype (Fig. 1A, B) is remarkable if one considers the estimate that about 50% of all NPCs are installed through interphase insertion (Doucet et al., 2010).

Based on our identification of MLF2 as a bleb-specific marker (Fig. 2), we developed a live cell imaging platform to show that bleb formation occurs synchronously within a narrow window of time immediately after NE reformation following open mitosis (Fig. 5A, Supplemental Video 1). Both the speed and synchrony of blebbing were entirely unexpected since we assumed a much broader, “stochastic” emergence of blebs based on experiments with fixed cells (Fig. 1D-F). Furthermore, it is noteworthy that a high degree of specificity exists for the luminal content. Model substrates of nuclear transport and ribosomes (some of the major nuclear export cargo) do not accumulate in these blebs (Fig. 3) (Laudermilch et al., 2016). This argues against the formal possibility that blebs merely occur upon the packaging of “random” nuclear export cargo. The question arises, however, as to whether K48-Ub or MLF2 play a role in NPC biogenesis or whether they are merely sequestered in blebs. We did not observe a major role for K48-Ub conjugation (Fig. 6C), and our preliminary MLF2 silencing approach did not suggest a critical role for MLF2 in bleb formation (Fig. S2D and E). Whether this is due to a possible genetic redundancy with the MLF2 homolog MLF1 remains to be seen. An alternative possibility is that the sequestration of MLF2 into blebs detrimentally affects the normal function of this protein, which is presently poorly understood (Banerjee et al., 2017; Kuefer et al., 1996).

Most importantly, our study firmly link Torsins to the process of interphase nuclear pore biogenesis. Apart from the aforementioned kinetics of bleb formation, this functional assignment is supported by the following observations: (i) a reduction in the number of mature pores (Fig. 1B), (ii) an underrepresentation of the late NPC assembly marker Nup358 from NEs of 4TorKO cells (Fig. 7) and (iii) the strict requirement of POM121–a transmembrane Nup essential for interphase assembly–for bleb formation (Fig. 8). In our model, NE blebbing during interphase NPC biogenesis serves to bring the INM within a fusogenic distance of the ONM (Fig. 9B). In this context, it might be useful to directly compare EM tomograms representing NE blebs in 4TorKO cells with those observed during interphase NPC biogenesis in unperturbed cells. The latter are somewhat flatter and dome shaped (Otsuka et al., 2016) while larger membrane herniations of about 200-250 nm are seen in 4TorKO cells (Laudermilch et al., 2016) (for a diagrammatic comparison, see Fig. 9A and B).

**Figure 9:**
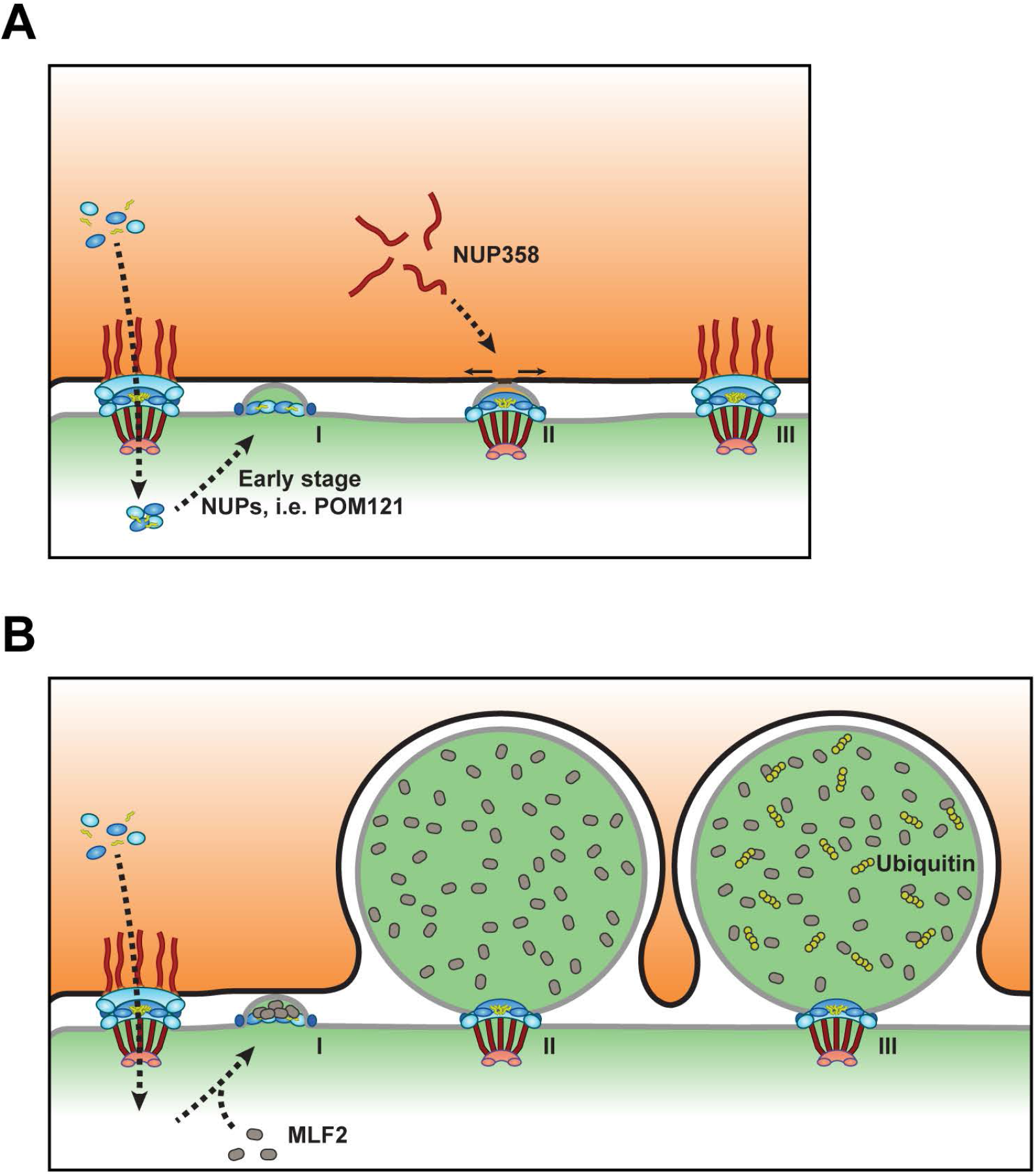
Models of normal interphase NPC assembly and defective NPC biogenesis resulting from Torsin manipulation. (A) Following mitosis, NPC assembly occurs via an inside-out evagination of the inner nuclear membrane (INM) in a process that requires the recruitment of POM121. Nuclear ring components and NPC subcomplexes are shuttled through mature pores previously assembled through a post-mitotic insertion mechanism. As pore intermediates mature, Nups that presumably deform the membrane evagination are added in a process that drives the growth of the complex both laterally and towards the outer nuclear membrane (ONM) (Intermediate I). Following a membrane fusion event, late stage and cytoplasmic Nups like Nup358 are added (Intermediate II) and the pore experiences significant architectural rearrangements ultimately giving rise to a complete NPC (III). (B) The formation of NE aberrations resulting from defective interphase assembly of NPCs upon Torsin manipulation or depletion. Interphase assembly begins in a POM121-dependent manner and pore intermediates mature to a stage that precedes membrane fusion (intermediate I). MLF2 is recruited early in the biogenesis of NE herniations, likely during the recruitment of early stage Nups, and is enriched in the lumen of the mature herniation. The INM proximal to the ONM expands to give rise to an omega-shaped herniation of the NE (Intermediate II). Specific protein components of the herniations including MLF2 are then labeled with K48 ubiquitin chains (Intermediate III). These structures are static through interphase and are turned over at the onset of mitosis.

How can we interpret this difference? We propose that in Torsin-deficient cells, the otherwise dynamically linked processes of INM deformation and INM/ONM fusion during NPC biogenesis is stalled before or at the step of INM/ONM fusion. As a first possibility, a specific NPC component might require Torsin for folding or trafficking, and the observed effects are indirect and result from a stalling since a specific required component is absent or misfolded. Based on our observation that POM121 is required for bleb formation (Fig. 8), we deduce that this component would likely have to be recruited downstream of POM121 in the assembly pathway. This interpretation would be consistent with previous observations of Ub conjugation in blebs. If we assume that one or several critical components important for NPC formation are misfolded due to the absence of Torsins and consequently ubiquitylated, its mere deubiquitylation would not be expected to restore protein function. As speculated before (Chase et al., 2017a; King and Lusk, 2016; Otsuka and Ellenberg, 2018), a second possibility is that Torsins might act on fusogenic components responsible for INM/ONM fusion. Since the fusogenic machinery remains to be identified, it is currently impossible to test this directly. Finally, Torsins have recently been linked to lipid metabolism (Grillet et al., 2016; Shin et al., 2019). We did not observe major changes in the lipid profile in 4TorKO cells vs. WT cells (Laudermilch et al., 2016), but we cannot exclude that local, NE-specific changes in lipid composition exist which might affect the fusogenic properties of NE membranes. It might be interesting to employ lipid-specific probes to scrutinize lipid composition of NEs and its possible perturbation in Torsin-deficient cells in the future.

Regarding the broader physiological implications of NE blebbing, it would be interesting to compare naturally occurring instances where blebbing phenotypes closely mirroring our observations have been documented (Thaller and Lusk, 2018). These include NE blebs during the neuromuscular junction in *D. melanogaster* (Jokhi et al., 2013; Speese et al., 2012), as well as NE blebbing that appears to be an evolutionary conserved process in zygotes and early embryos. In the latter case, blebs with necks of dimensions similar to NPCs were observed, although they have not been linked to NPC biogenesis (Szollosi and Szollosi, 1988). It will be interesting to test whether these can be decorated with Mab414 antibodies, and if these structures contain MLF2. It is tempting to speculate that several of these observations can in fact be connected to NPC biogenesis.

Regardless of these questions, we interpret our results to firmly link Torsin ATPases to the process of interphase NPC biogenesis. Our findings have distinct implications for our understanding of movement disorders caused by Torsin dysfunction in neurons. Neuronal cells display a low mitotic index and are thus expected to be particularly vulnerable since these are more dependent on the interphase assembly pathway than dividing cells, which can utilize the alternate post-mitotic insertion pathway. Additionally, neurons are heavily reliant on TorsinA due to a window in neurogenesis during which TorsinA is the dominantly expressed Torsin relative to its homologs (Kim et al., 2010; Tanabe et al., 2016).Thus, our results, along with these data, suggest that defects in NPC biogenesis add to DYT1 dystonia’s disease etiology.

## Materials and Methods

### Tissue culture

Torsin-deficient HeLa cells and their parental WT cell line were cultured as previously described (Laudermilch et al., 2016). Briefly, cells were cultured in Dulbecco’s Modified Eagle’s Medium supplemented with 10% fetal bovine serum (FBS) (Thermo Fischer Scientific) and 100 units mL^-1^ of penicillin-streptomycin (Thermo Fischer Scientific). Cells were routinely checked for mycoplasma and determined to be free of contamination through the absence of extranuclear Hoechst 33342 (Life Technologies) staining.

### Plasmids constructs

The sequence encoding MLF2 was amplified by standard PCR procedures from cDNA (Dharmacon; Accession # BC000898) and subcloned into either a pcDNA3.1^+^ vector (MLF2-HA and MLF2-Flag) or a pEGFP-N1 vector (MLF2-GFP). MLF2-M48 fusion variants were constructed through standard Gibson assembly procedures. Plasmids containing the WT and catalytic mutant variant of M48 were a gift from Hidde L. Ploegh (Whitehead Institute for Biomedical Research) (Schlieker et al., 2007). The plasmid containing Sec61β was gifted from Shirin Bahmanyar (Yale University). Sec61β was subcloned into a modified pEGFP-C1 vector in which EGFP was replaced with mScarlet (pmScarlet-C1). Both MLF2-GFP and mScarlet-Sec61β were subcloned into the pRetroX-Tight-Pur-GOI vector (Takara Bio). NES-GFP was custom synthesized as a 789 bp gBlock gene fragment (Integrated DNA Technologies) with an N-terminal nuclear export sequence (Cardarelli et al., 2012). NLS-GFP was gifted from Anton Bennett (Yale School of Medicine). WT RAN was derived from pmCherry-C1-RanQ69L (Addgene: 30309) by reverting the point mutation through site directed mutagenesis. WT RAN was subcloned with an N-terminal HA tag into pcDNA3.1^+^.

### Generation of HeLa stable cell lines

To generate a 4TorKO cell line stably expressing MLF2-GFP and mScarlet-Sec61β, we employed the Retro-X Tet-On advanced inducible expression system (Takara Bio) following the manufacturer’s protocol. For the production of retrovirus, low-passage 293T cells were transfected with 2 µg MMLV gag/pol, 1 µg viral envelope protein VSV-G, and 6 µg of either pRetroX-Tight-Pur-MLF2-GFP, pRetroX-Tight-Pur-mScarlet-Sec61β, or pRetroX-Tet-On using X-tremeGENE 9 (Roche).

Supernatants containing retroviruses were collected 72 h post-transfection, filtered via a 0.45-μm filter unit, and stored at −80°C. 4TorKO cells were seeded in 6-well plates 24 hrs prior to transduction. The next day, media was replaced with complete growth media supplemented with 4 μg mL^-1^ polybrene (Sigma-Aldrich) and 100 µL of the respective retroviruses were added dropwise to the wells. Media was replaced 24-hours post transfection to fresh complete media containing 1 µg mL^-1^ puromycin (Sigma-Aldrich) and 800 μg mL^-1^ Geneticin (Thermo Fisher Scientific). Antibiotic selection was performed for 7 days. Cells positive for both GFP and mScarlet signal under the Dox-inducible promoter were isolated through fluorescence activated cell sorting (FACS). FACS was performed at the Yale University Flow Cytometry Facility using aFACS Aria III sorter (BD Biosciences).

To establish WT and 4TorKO cells stably expressing MLF2 with an endogenous C-terminal 3xHA tag, we utilized a CRISPR/Cas12a system for PCR tagging genes (Fueller et al., 2018). Oligo sequences targeting MLF2 at its endogenous locus were generated from an open access tool (www.pcr-tagging.com). Sequences for the PCR tagging oligo primers are M1: 5’-GCTGGGGGACGAAGGGCGGAGGGGCCTCCCCGCCTGGCCATCCAGGGACCTGAGGACTCCCCTTCCCGACAGTCCCGCCGCTATGACTGGTCAGGTGGAGGAGGTAGTG-3’ and M2: 5’-CACCCCACCCTCCTTACTCCTGATACTTACAAGAGAGGCTGAGGGCCCGGGGCCCAAAAAAGGCCCGGGGCCCTCACCAGTATCTACAAGAGTAGAAATTAGCTAGCTGCATC GGTACC-3’ (Integrated DNA Technologies). pMaCTag-P28 (Addgene: 120039) was used as PCR template. Cells were transfected with 1 ug total DNA (0.5 ug PCR product and 0.5 ug AsCpf1_TATV Cas12 [Addgene: 89354]) using Xtreme-GENE 9 following manufacturer instructions. Media was replaced 24hr post transfection to fresh complete media containing 1 µg/mL Puromycin (Sigma-Aldrich). Antibiotic selection was performed for seven days. Following selection, cells derived from individual colonies were screened for MLF2-3xHA fusion protein by immunoblot, and colonies with HA signal were propagated and saved.

### Cell synchronization

Cells were synchronized with a double thymidine block (Bostock et al., 1971). Cells were incubated in complete growth media supplemented with 2.5 mM thymidine (Sigma-Aldrich) for 18 h. Cells were released from the thymidine block by washing with Dulbecco’s phosphate-buffered saline (DPBS) (Thermo Fischer Scientific), and replacing the media with fresh complete growth media. Cells were incubated for 9 h at which point a second round of 2.5 mM thymidine treatment was administered. Following a 16-hour incubation, cells were again washed and incubated in complete growth media without thymidine. This final media replacement was designated as T = 0, and time points were then collected afterwards as indicated in the text.

### siRNA knockdown and RT-qPCR validation

siRNA knockdown was done using Lipofectamine RNAimax (Life Technologies). The Lipofectamine reagent was diluted in Opti-MEM reduced serum medium (Thermo Fischer Scientific) for 15 min, followed by the addition of the appropriate concentration of siRNA and incubated further for 15 min. This solution was then added dropwise to cells and allowed to incubate overnight. Media was then replaced with fresh antibiotic-free media. Cells were harvested for qPCR or immunofluorescence (IF) 48 h (siNT, siELYS, and siMLF2) or 72 h (siPom121) post-transfection. Forward siRNA sequences 5’-CAGUGGCAGUGGACAAUUCA[dT][dT]-3’ (Sigma) and 5’-UCGUGGAAAGUUUGCUGCAGGGAAA[dT][dT]-3’ (Sigma) (Doucet et al., 2010) were used for POM121 and ELYS, respectively, while SMARTpool siRNA was utilized to target MLF2 (Dharmacon). Following treatment, cells were either fixed for IF analysis or total RNA was extracted for qPCR following previously described methods (Tsai et al., 2016). In short, 100 ng of RNA was transcribed into cDNA using SuperScript II reverse transcriptase (ThermoFisher Scientific) with random hexamer primers (Invitrogen). qPCR was performed using iQ SYBR Green mix and executed on a CFX Real-Time PCR 639 Detection System (Bio-Rad). The ΔΔCt values for each sample were calculated from the subtraction of an internal control value (GAPDH) and results were normalized to the siNT control. Primer sequences (5’-3’) utilized for qPCR were as follows: GAPDH (Forward: CGACCGGAGTCAACGGATTTGGTCG; Reverse: GGCAACAATATCCACTTTACCAGA), ELYS (Forward: CCAATTTCTGACAGCCCTCCTGA; Reverse: AGATTCCTAGCCTCTTCTCCTGAA), POM121 (Forward: CCTTCAGCCAGTCCCTGCAC; Reverse: GAGGGTGCTGCCAAAACCAC), and MLF2 (Forward: GGACTCCCCTTCCCGACAGT; Reverse: GCCTCTCAGCCTGTACAAGAG) (Integrated DNA Technologies).

### Nuclear envelope isolation and K48 ubiquitin immunoprecipitation

The isolation of NE membranes was modified from previously described methods (Emig et al., 1995; Tsai et al., 2019). Briefly, WT and 4TorKO cells were collected from 5 15-cm plates and centrifuged at 500 x g for 5 minutes at 4°C. Cells were resuspended in cold PBS and 100 µL of cells were set aside for WCL input controls. Cells were again centrifuged and resuspended in 5 mL of cold Buffer A (10 mM HEPES, pH 7.4, 250 mM sucrose, 2 mM MgCl_2_) supplemented with 1 mM phenylmethylsulfonyl fluoride (PMSF) and incubate on ice for 10 min. Cells were then homogenized by passing through a 25G needle 5 times. Homogenates were transferred to the top of 10mL STM 0.9 buffer (50 mM Tris, pH 7.4, 0.9 M sucrose, 5 mM MgCl_2_) and sedimented at 1,000 x g for 10 min. Pellets containing crude nuclear fractions were resuspend in 5 mL STM 1.6 buffer (50 mM Tris, pH 7.4, 1.6 M sucrose, 5 mM MgCl_2_,1 mM PMSF). Suspensions were underlayed with 1 mL STM 2.1 buffer (50 mM Tris, pH 7.4, 2.1 M sucrose, 5 mM MgCl_2_) and 4 mL STM 0.8 buffer (50 mM Tris, pH 7.4, 0.8 M sucrose, 5 mM MgCl_2_) was added as the top layer. Pure nuclear fractions were sedimented by ultracentrifugation at 28,500 rpm (rotor SW41) for 65 min. Nuclear pellets were washed once in 1 mL TP buffer (10 mM Tris, pH 8.0, 10 mM Na_2_HPO_4_, 5 mM MgCl_2_) and sedimented at 1000 x g for 10 min at 4°C. Nuclear pellets were resuspended in 0.5 mL TP buffer supplemented with heparin (7.2 mg / 24 ml buffer), 1 µL benzonase, and 2 mM NEM and rocked at 4°C for 2 h. Samples were centrifuged at 15,000 x g for 10 min at 4°C. Supernatants containing NP fractions were saved and pellets containing NE fractions were solubilize in 1 mL solubilization buffer (50 mM Tris, pH 7.5, 5 mM MgCl_2_, 150 mM NaCl and 2% digitonin, 1mM PMSF, 2mM NEM) on ice for 30 min. Samples were centrifuged at 15,000 x g for 10 min at 4°C and the supernatant was transferred to a clean microcentrifuge tube. A 15µL aliquot was set aside to assess the quality of the fractionation.

Equal protein concentrations from WT and Torsin-deficient cells were immunoprecipitated. Immunoprecipitation was performed with 5 µL anti-K48 ubiquitin (AB_11213655, Millipore) conjugated to protein A Dynabeads for 3 h at 4°C. Beads were washed three times with wash buffer (50 mM Tris, pH 7.5, 150 mM NaCl, 5 mM MgCl_2_, 0.1% digitonin) and proteins were eluted by heating to 65°C for 5 min in 30 µL SDS loading buffer. Eluates were subjected to SDS-PAGE on a Mini-PROTEAN precast gel (Bio-Rad). Immunoblotting was performed to assess the quality of the fractionation protocol with anti-SUN2 antibody (AB_1977547, Millipore), and anti-HA antibody (AB_390919, Roche) at a 1:5000 and a 1:4000 dilution, respectively. Gels for mass spectrometry (MS) analysis were stained with SimplyBlue Safe Stain (Thermo Fischer Scientific) and gel samples were sent to the Yale NHLBI Proteomics Center for LC-MS/MS. MS proteomic data was analyzed with Scaffold (Proteome Software Inc., Portland, Oregon).

### Transmission electron microscopy

Electron microscopy was performed at the Center for Cellular and Molecular Imaging, Yale School of Medicine with a previously described workflow (Laudermilch et al., 2016). Briefly, cells were fixed for 1 h in 2.5% glutaraldehyde in 0.1 M sodium cacodylate buffer, pH 7.4. Following a brief rinse, cells were scrapped in 1% gelatin and centrifuged in a 2% agar solution. Chilled cell blocks were processed with osmium and thiocarbohydrazide-osmium liganding as previously described (West et al., 2010), and samples were embedded in Durcupan ACM resin (Electron Microscopy Science). Polymerization was performed by incubating samples at 60°C overnight. These blocks were cut into 60 nm sections with a Leica UltraCut UC7, and stained with 2% uranyl acetate and lead citrate on Formvar/carbon-coated grids.

Samples for immunoelectron microscopy were processed as described above with some modifications. Samples were fixed through high-pressure freezing (Leica EM HPM100) and freeze substitution (Leica AFS) at 2000 PSI. Freeze substitution was performed by incubating samples in 0.1% uranyl acetate/acetone solution for 50 h at −90°C. Following treatment, samples were washed in acetone, infiltrated in Lowicryl HM20 resin (Electron Microscopy Science) for 10 h at −45°C, transferred to gelatin capsules, and hardened via ultraviolet light exposure at - 45°C. Blocks were subsequently sectioned and placed on Formvar/carbon-coated nickel grids for immunolabeling. Untreated aldehyde groups were quenched by incubating grids in 0.1 M ammonium chloride prior to immunolabeling procedure. Samples were subsequently blocked in 1% fish-skin gelatin in PBS and grids were incubated in a 1:50 dilution of anti-GFP antibody (AB_390913, Roche) and labeled with 10 nm Protein A-gold particles (Utrecht Medical Center). Grids were fixed using 1% glutaraldehyde, washed with PBS, dried, and stained using 2% uranyl acetate and lead citrate.

Both conventional EM and immunogold labeled samples were visualized with an FEI Tecnai Biotwin TEM at 80Kv and pictures were taken with Morada CCD and iTEM (Olympus) software.

### Immunofluorescence for wide field fluorescence and confocal microscopy

IF staining was done as previously described (Laudermilch et al., 2016; Rose et al., 2014; Tsai et al., 2016). Briefly, cells were cultured on coverslips (VWR), fixed with 4% paraformaldehyde (PFA) in PBS, and permeabilized for 10 mins with 0.1% Triton-X 100 (Sigma-Aldrich) in PBS. Cells were blocked with 4% bovine serum albumin (BSA) (Sigma-Aldrich) in PBS for 10 mins and incubated with the appropriate antibodies for 45 mins. The following antibodies were used at a 1:500 concentration: anti-K48 ubiquitin (AB_11213655, Millipore), anti-HA (AB_390919, Roche), and anti-Flag (AB_259529, Sigma-Aldrich). After five washes, cells were blocked with 4% BSA and incubated with secondary antibodies conjugated to Alexa-Fluor™ 488 (Life Technologies) or Alexa-Fluor™ 568 (Life Technologies) for 45 mins. Cells were washed with PBS, incubated in Hoechst 33342 (Life Technologies) for 5 mins, and washed in PBS wash before being mounted onto slides using Fluoromount-G (Southern Biotech).

Standard wide field images were obtained with a Zeiss Axio Observer D1 microscope using a 63x oil immersion objective. Confocal images were taken with an LSM 880 laser scanning confocal microscope (Zeiss) using a C Plan-Apochromat 63x/1.40 Oil DIC M27 objective.

### Immunofluorescence for 3D-SIM

Cells grown on coverslips were fixed in 4% PFA for 10 mins and washed with PBS. Blocking and antibody dilutions were carried out in 3% BSA in PBS with 0.1% Triton-X 100. After 1 h blocking, cells were incubated with an anti-Nup358 antibody (kind gift of Gunter Blobel/Elias Coutavas) for 1 h at room temperature (RT). Cells were washed with three times with PBS for 5 min each and incubated with Alexa-Fluor™ 568 goat anti-rabbit (Thermo Fisher Scientific) for 1 h at RT. Cells were washed as described above and then incubated with the Mab414 antibody (1:500, Abcam) for 1 h. Cells were washed as described above and then incubated with Alexa-Fluor™ 488 goat anti-mouse (Thermo Fisher Scientific) for 1 h at RT. Cells were washed and mounted using Fluoromount-G™ (Electron Microscopy Sciences) before imaging.

### 3D-SIM

3D-SIM imaging was performed on a DeltaVision OMX V3 system (GE Healthcare Life Sciences) equipped with a U-PLANAPO 60X/1.42 PSF oil immersion objective lens (Olympus, Center Valley, PA), CoolSNAP HQ2 CCD camera with a pixel size of 0.080 µm (Photometrics) and 488 nm, 561 nm, and 642 nm solid-state lasers (Coherent and MPB communications). Image stacks were acquired in 0.125 µm increments in the z-axis in sequential imaging mode. Samples were illuminated by a coherent scrambled laser light source first passed through a diffraction grating to generate the structured illumination by interference of light orders in the image plane to create a 3D sinusoidal pattern, with lateral stripes approximately 0.270 µm apart. The pattern was shifted laterally through five phases and through three angular rotations of 60° for each Z-section, separated by 0.125 µm. Exposure times were typically between 25 and 150 ms, and the power of each laser was adjusted to achieve optimal fluorescence intensities between 2,000 and 4,000 in a raw image of 16-bit dynamic range, at the lowest possible laser power to minimize photo bleaching. Color channels were carefully aligned using alignment parameters from control measurements with 0.5 µm diameter multi-spectral fluorescent beads (Thermo Fisher Scientific).

The 3D-SIM images were subjected to SIM reconstruction and image processing using the SoftWoRx 3.7 imaging software package (GE Healthcare Life Sciences). The channels were then aligned in x, y, and rotationally using predetermined shifts as measured using a target lens and the SoftWoRx alignment tool (GE Healthcare Life Sciences).

### Live cell lattice light sheet microscopy

4TorKO cells expressing mScarlet-Sec61β and MLF2-GFP under a Dox-inducible promoter were cultured on 5-mm diameter coverslips (Warner Instruments) in complete media 24 h prior to cell cycle synchronization. Media was supplemented with 0.5 µg mL^-1^ doxycycline 24 h prior to imaging. Coverslips were mounted and fixed to a temperature-controlled imaging chamber containing Leibovitz’s L-15 medium (Life Technologies, 11415064) supplemented with 10% FBS equilibrated to 37°C. MLF2-GFP and mScarlet-Sec61β were excited using a 488-nm laser and a 560-nm laser, respectively, with inner and outer numeric apertures of 0.325 and 0.4, respectively, and 5 ms exposure times. Imaging data was acquired with a sCMOS camera (Hamamatsu Orca Flash 4.0 v3). Metaphase cells were identified, and three-dimensional data sets were recorded with 20 s intervals between time points. All LLSM data was deconvolved with the Janelia open source software cudaDeconv (Janelia).

### Image processing and data analysis

All indirect IF, confocal, and 3D SIM images were processed for figures and analyzed with FIJI software (Schindelin et al., 2012). In addition to FIJI software, LLSM images were also processed with FluoRender 2.19 (Scientific Computing and Imaging Institute, Salt Lake City, UT).

Quantifications of blebs were performed as previously described (Rose et al., 2014). Briefly, cells were randomly selected using the Hoechst channel and were imaged for K48 ubiquitin staining. Nuclei were outlined manually, and the number of foci per nucleus was determined by the “Find Maxima” function in FIJI with a noise tolerance of 10. A threshold was set by comparison to wild type nuclei, and cells above the threshold were determined to contain K48-Ub foci. Statistical analysis was performed in GraphPad Prism.

Line scan analyses were performed in FIJI. The nuclear periphery of a given region was traced with the segmented line selection tool and the same trace was superimposed on subsequent channels for consistency through the “Restore Selection” function. A plot of the intensity profile for the selected region was generated through the “Plot Profile” function and all data was exported and graphed using GraphPad Prism.

Quantifications for the increase in fluorescence intensities of MLF2-GFP foci were performed in FIJI. Individual foci were traced with the freehand selection tool and the average pixel intensities per foci area were determined through the measure function. Average fluorescence intensities are represented as relative intensities units (RIU) and the RIU values of each foci for a given time point are normalized by subtracting the background intensity of the same area of a respective focus. Data was exported and graphed using GraphPad Prism

## Contributions

A.J. Rampello, E. Laudermilch, C. Zhao, N. Vishnoi, L. Shao, S.M. Prophet, C.P. Lusk, and C. Schlieker conceptualized and designed experiments in the text. A.J. Rampello, E. Laudermilch, C. Zhao, N. Vishnoi, L. Shao, and S.M. Prophet performed experiments. A.J. Rampello, E. Laudermilch, C. Zhao, N. Vishnoi, L. Shao, S.M. Prophet, C.P. Lusk, and C. Schlieker analyzed and interpreted data. A.J. Rampello, S.M. Prophet, and C. Schlieker wrote the original manuscript. A.J. Rampello, E. Laudermilch, C. Zhao, N. Vishnoi, L. Shao, S.M. Prophet, C.P. Lusk, and C. Schlieker revised and edited the manuscript.

## Acknowledgements

This work is supported by NIH R01GM114401 (C.S.), NIH 5T32GM007223-44 (S.M.P.), NIH GM105672 (C.P.L.) and the Dystonia Medical Research Foundation (C.S., A.J.R. and C.P.L.). We thank Joerg Bewersdorf and members of his laboratory for continued support and Felix E. Rivera-Molina for help with 3D-SIM.

---

**Supplemental Figure 1:**
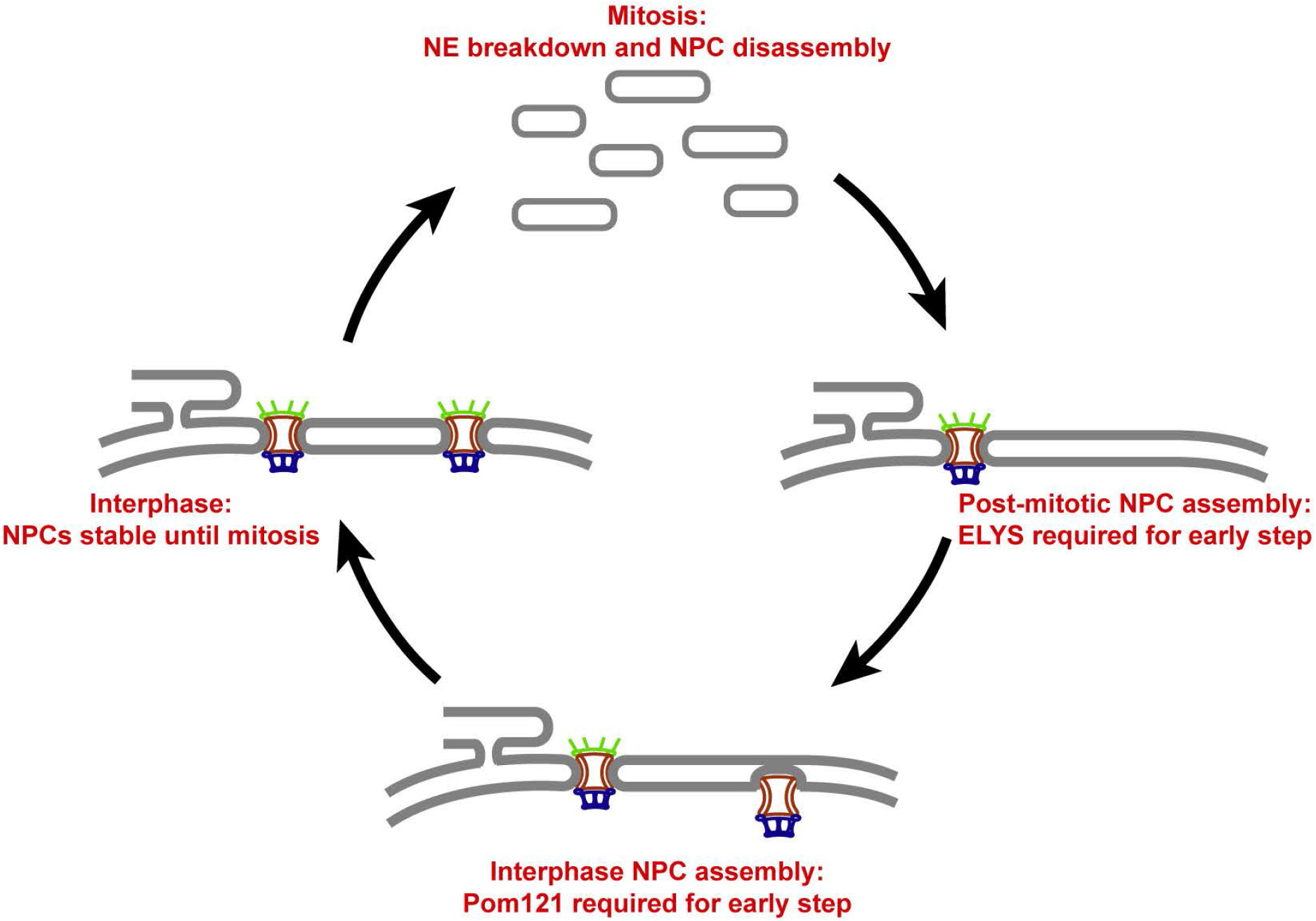
Distinct stages of the NPC lifecycle. Graphical depiction of various stages of the NPC lifecycle. During mitosis, NPCs are disassembled and the nuclear envelope retreats to the ER (top panel). NPCs can then assemble as cells exit mitosis via the post-mitotic pathway (right panel) or during interphase (bottom panel). Pores formed by both pathways are likely stable until mitosis (left panel).

**Supplemental Figure 2:**
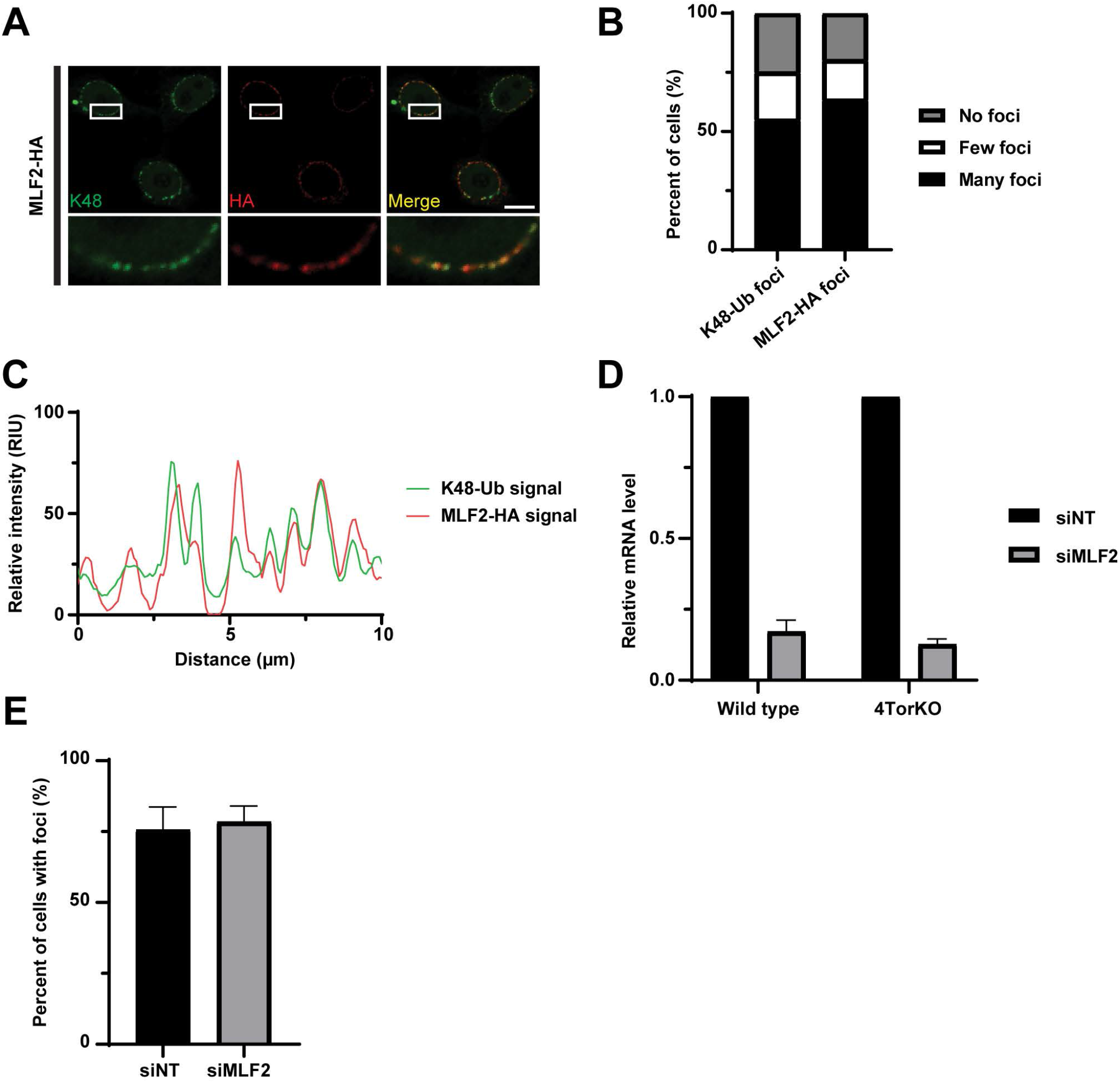
MLF2 localization correlates with K48 ubiquitin enrichment at the nuclear periphery. (A) Confocal microscopy images demonstrating the colocalization between MLF2-HA and K48-Ub at the nuclear periphery of 4TorKO cells. Representative scale bar is 10 µm. (B) Graphical representation illustrating the percent of 4TorKO cells transfected with MLF2-HA that display K48-Ub foci or MLF2-HA foci at the nuclear periphery. Data shown are the mean of three independent experiments with at least 75 cells each. (C) Line-scan profiles of K48-Ub signal (green) and MLF2-HA signal (red) at the nuclear periphery of the cell from (A). Measurements were taken for the region represented in the inset. (D) The relative amount of MLF2 mRNA transcript following a 50 nM siRNA treatment was analyzed by RT-qPCR. Relative mRNA levels were normalized to the siNT control. (E) The percent of 4TorKO cells with K48-Ub foci upon treatment with 50 nM siNT or siMLF2. The values are means of three independent replicates (n = 3) and error bars indicate ± SD.

**Supplemental Figure 3:**
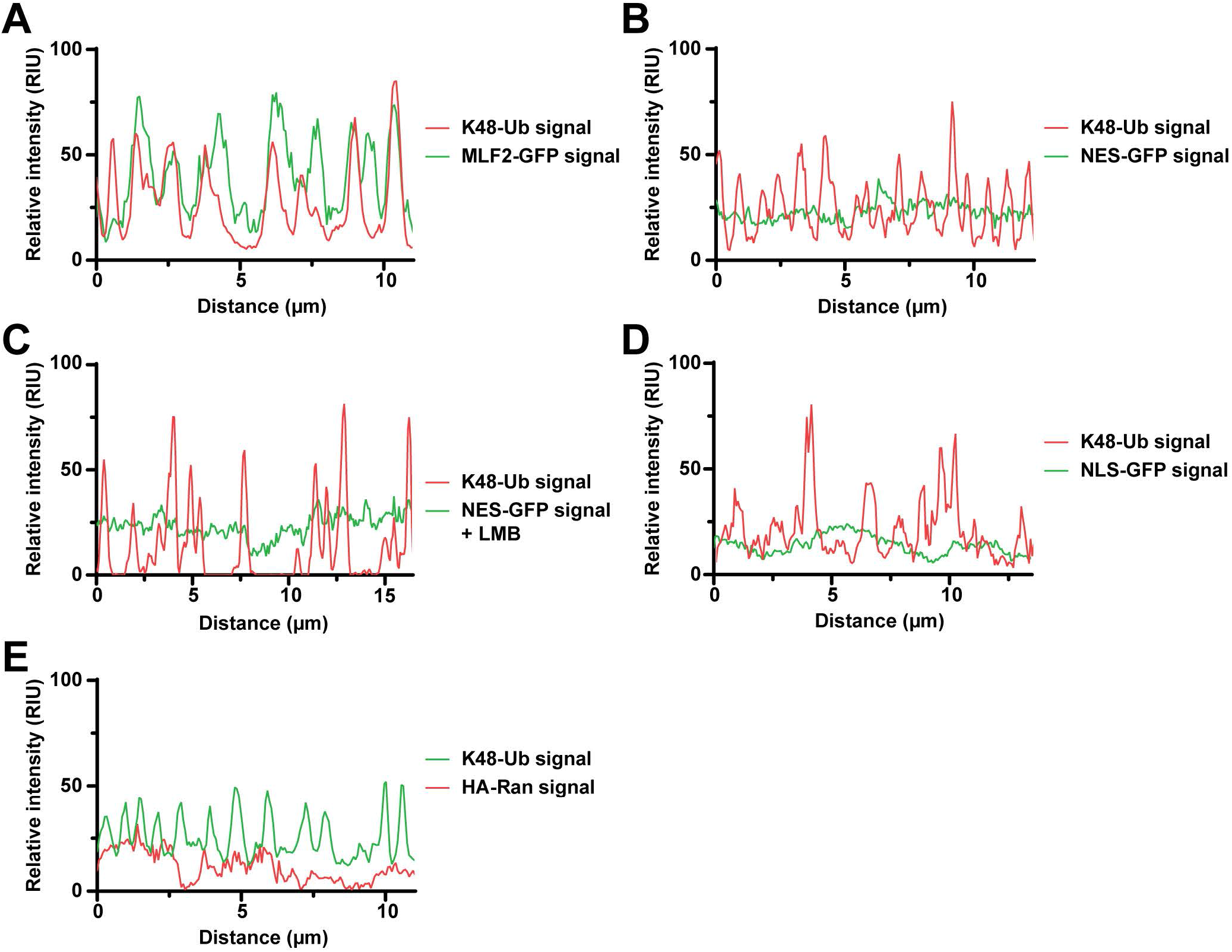
Signal intensities of MLF2-GFP, NES-GFP, NLS-GFP, and HA-RAN relative to K48 ubiquitin signal at the nuclear periphery. (A-D) Line-scan profiles of K48-Ub (red) and MLF2-GFP (green), NES-GFP (green), NES-GFP under a 10 ng mL^-1^ LMB treatment (green), NLS-GFP (green), and (E) K48-Ub (green) with HA-Ran (red) at the nuclear periphery. Measurements were taken for the region represented in the inset images from Figure 3A-D.

**Table S1:** List of proteins identified by mass spectrometry after subcellular fractionation and K48-Ub immunoprecipitation. Proteins shown are the top 20 candidate proteins scored by highest percent coverage (%) in 4TorKO cell IP.

|  | Quantitative Value<br>(Normalized Total Spectra) |  | Total Unique Peptide Count |  | Percent coverage (%) |  |
| --- | --- | --- | --- | --- | --- | --- |
| Protein Name | 4TorKO IP | WT IP | 4TorKO IP | WT IP | 4TorKO IP | WT IP |
| UBA52 | 49.16 | 35.03 | 11 | 11 | 50.00 | 50.00 |
| ACTC1 | 33.09 | 27.60 | 25 | 16 | 47.20 | 44.30 |
| HSPA1A | 29.30 | 7.43 | 23 | 6 | 36.20 | 11.90 |
| **MLF2** | **7.56** | **2.12** | **8** | **2** | **35.90** | **12.10** |
| HSPA8 | 48.21 | 25.48 | 27 | 20 | 34.80 | 33.60 |
| VCP | 29.30 | 31.84 | 26 | 29 | 34.60 | 38.20 |
| EMD | 8.51 | 1.06 | 8 | 1 | 33.50 | 4.72 |
| UBAC2 | 8.51 | 3.18 | 8 | 3 | 29.90 | 13.70 |
| DNAJB2 | 8.51 | 1.06 | 6 | 1 | 27.80 | 3.09 |
| DNAJB6 | 7.56 | 2.12 | 5 | 2 | 23.60 | 8.97 |
| ERLIN2 | 7.56 | 11.68 | 6 | 6 | 22.40 | 22.40 |
| ACTBL2 | 8.51 | 13.80 | 7 | 8 | 20.50 | 26.30 |
| SARAF | 8.51 | 0.00 | 5 | 0 | 19.20 | 0.00 |
| VDAC1 | 5.67 | 9.55 | 5 | 7 | 18.70 | 33.60 |
| APOA1 | 3.78 | 10.61 | 4 | 9 | 18.70 | 34.50 |
| HSPA9 | 10.40 | 8.49 | 9 | 8 | 18.00 | 16.20 |
| CANX | 6.62 | 11.68 | 7 | 7 | 15.50 | 16.20 |
| DNAJB12 | 5.67 | 0.00 | 5 | 0 | 13.60 | 0.00 |
| PDE12 | 5.67 | 3.18 | 6 | 3 | 13.50 | 6.24 |
| KCNG1 | 9.45 | 0.00 | 7 | 0 | 10.50 | 0.00 |

<u>Supplemental Video 1:</u> Formation of nuclear envelope (NE) blebs visualized by lattice light sheet microscopy. A maximum intensity projection of a 3D image stack (z-axis) depicting 4TorKO cells expressing mScarlet-Sec61B and MLF2-GFP. Video captures anaphase onset (0 s) and follows cells through cytokinesis. MLF2-GFP foci form rapidly in early G1 phase (∼700 s). Foci formation is followed by a fast growth phase (∼700 s - on). Video corresponds to Figure 5A, C, D, and E.

<u>Supplemental Video 2:</u> MLF2-GFP foci localize to NE membranes. Z-scan through recently divided 4TorKO cells expressing mScarlet-Sec61B and MLF2-GFP. The majority of MLF2-GFP foci form on the NE and its membranous protrusions extending into the nucleus. Video corresponds to Figure 5B.

